# EzMechanism: An Automated Tool to Propose Catalytic Mechanisms of Enzyme Reactions

**DOI:** 10.1101/2022.09.05.506575

**Authors:** Antonio J. M. Ribeiro, Ioannis G. Riziotis, Jonathan D. Tyzack, Neera Borkakoti, Janet M. Thornton

## Abstract

A rich literature dedicated to understanding the reaction mechanisms of hundreds of enzymes has emerged over time from the works of experimental and computational researchers. This body of information can now be the starting point for an entirely novel approach to studying enzyme mechanisms using knowledge-based prediction methods. Here, we present such a method, EzMechanism, (pronounced as “Easy Mechanism”) which is able to automatically generate mechanism proposals for a given active site. It works by searching the chemical reaction space available to the enzyme using a set of newly created biocatalytic rules based on knowledge from the literature. EzMechanism aims to complement existing methods for studying enzyme mechanisms by facilitating and improving the hypotheses generating step. We show that EzMechanism works by validating it against 56 enzymes with a known mechanism and identify the limited coverage of the current ruleset as the main target for further improvement.

## Introduction

Enzymes are proteins that accelerate the chemical reactions necessary for life. These catalytic macromolecules are abundant in all genomes (22% of the coded proteins in humans and 40% in *E. coli*, for example^1^) and are widely studied. Of particular interest is understanding the enzymes’ reaction mechanisms: the sequence of events in the active site, such as the formation and cleavage of bonds, that explain how the substrate is transformed into its products. Enzyme mechanisms are crucial to understand enzyme function and evolution. This knowledge opens the door to a host of green-chemistry and biotechnological applications, from the modulation of enzyme function through rational design^2^, to the prediction of the impact of enzyme variants in disease^3^, and the development of drugs targeted at the active site^4^.

Learning about new enzyme mechanisms is a complex task, requiring the application of diverse types of experimental and computational methods. Enzymatic reaction rates, as well as the rate dependence on pH, temperature, or chemical species like cofactors, can be inferred from kinetic assays^5^. Potential catalytic sites identified among highly conserved residues^6^ can be confirmed by mutagenesis studies^7^. Spectroscopy data, such as EPR (Electron Paramagnetic Resonance) for metals and radical species^8^, or fluorescence for fluorescent intermediates^9^, may confirm or exclude the presence of certain molecular species along the reaction path. Three-dimensional structures^10^ provide information about the precise location of catalytic residues, substrates, and cofactors in the active site. Finally, computational chemistry methods such as QM/MM^11,12^, have been used to simulate reaction mechanisms with increasingly accurate models of the enzyme.

With the help of these methods, researchers have succeeded in building a rich literature about all aspects of enzyme function, including their reaction mechanisms. We have captured some of this information in the M-CSA (Mechanism and Catalytic Site Atlas)^13^, a manually curated database of enzyme mechanisms and catalytic sites that is freely available at www.ebi.ac.uk/thornton-srv/m-csa/. Among other data, M-CSA contains stepwise descriptions of the mechanistic path of 745 enzymes, stored as two-dimensional curly arrow diagrams (see Fig 1-A), denoting the movement of electrons as bonds are broken and formed during catalysis. This machine-readable representation of the mechanism of enzymes is unique to the M-CSA database and is the foundation of the present work.

**Figure 1.**
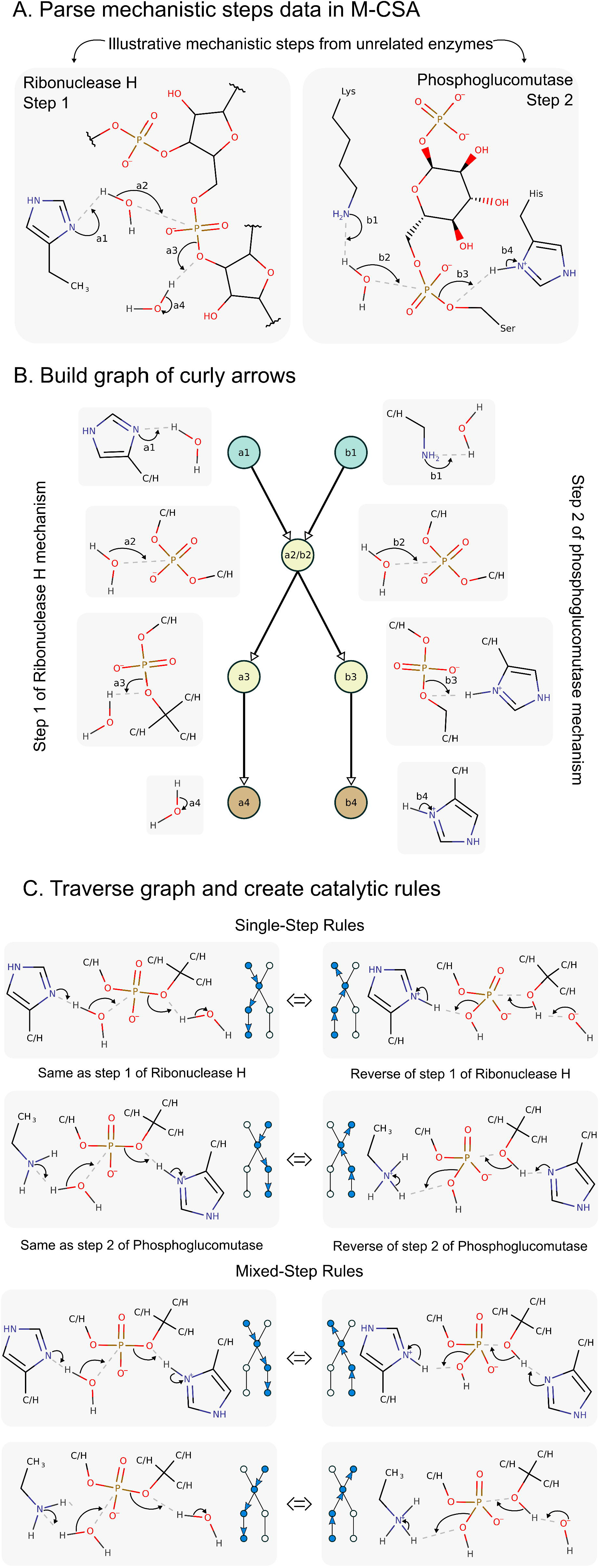
The process followed to generate the rules of biocatalysis as extracted from the M-CSA database, explained for two mechanistic steps. A – Example of two mechanistic steps from unrelated enzymes that share some chemical similarities. B – Graph of curly arrows showing the electron flow in the chemical groups seen in the exemplified steps. C – Single-step and mixed-step rules that can be created by traversing the curly arrow graph. The small version of the curly arrow graph besides each rule indicates how the graph must be traversed to generate that rule.

In this paper, we describe EzMechanism, a new tool that is able to suggest possible reaction mechanisms for a given enzyme active site. Our approach to develop EzMechanism was to first derive a set of “rules of biocatalysis” based on known mechanisms, as annotated in M-CSA. Then, we created a search algorithm that given an enzyme active site and overall reaction, uses these rules to explore the reaction space of the active site in order to find potential reaction mechanisms. A visualisation tool facilitates the inspection of these proposals, which can be compared to known mechanisms and prioritised based on geometrical (3D distances between atoms that form new bonds) or topological (such as number of steps) parameters.

We validated EzMechanism by running it against 56 M-CSA entries for which the mechanism is already known. For the purposes of this validation, we were particularly interested in assessing how well the prioritisation and search algorithms work, how comprehensive the catalytic rules are, and whether EzMechanism can find alternative mechanisms not previously considered in the literature.

EzMechanism proposals are generated in a matter of a minutes up to a few hours, depending on the complexity of the active site, and should be particularly useful for researchers in the initial stages of studying an enzyme mechanism, when different hypotheses for the mechanism are being considered. The search performed by the software is purely based on the chemical groups of the molecules in the active site which means that it can use information from related, evolutionary distant or even unrelated enzymes. EzMechanism hypotheses should then be tested experimentally or computationally using more sophisticated methods, such as QM/MM calculations.

EzMechanism is the first computational method that is able to come up with potential enzyme mechanisms for a given 3D structure of an active site (we discuss a related knowledge-based method^14^ developed independently in the online methods). We show that the software is efficient at searching the catalytic space and find productive paths between the reactants and products of enzyme reactions. The current version of EzMechanism is limited to non-radical reactions and our validation shows that in order to cover the complete enzymatic space, the created biocatalytic rules need to be improved and augmented. Together with tackling these limitations, future versions of the software will consider conformational variability and dynamics and use energy calculations to better guide the search.

In an ideal world, computers would be able to predict how enzymes work and the reactions they perform *ab initio*, based solely on their sequences. This would entail being able to predict their structure, what are the enzyme’s substrates and products (both native and potential new ones), as well as predicting the effect of mutations and conformation variability on the mechanism. As other computational methods in the fields of bioinformatics and computational chemistry keep developing (e.g. AlphaFold^15,16^, Rosetta, QM/MM^11,12^, molecular dynamics^17^, molecular docking^18^), we see EzMechanism as a crucial cog in this ideal vision of the future, as a way to codify and apply existing chemical knowledge to make predictions.

## Results

### The Rules of Biocatalysis

Enzymes use a diverse set of active sites to catalyse a large number of reactions, but the building units of these active sites are strikingly limited. Less than half of the 20 amino acids regularly play a direct role in the mechanism^6^, and the number of available cofactors is equally restricted^19^. This means that many enzymes will necessarily share components of their mechanisms, even if these enzymes are not evolutionarily related or catalyse different chemical reactions. For related enzymes the similarities will be even higher. The first step towards our goal was to identify these recurring “mechanistic components”, which we called the “rules of biocatalysis”. These rules codify chemical transformations that are possible when certain chemical groups are observed in the active site (typically there is one rule for each catalytic step) and can be written as simple chemical reaction equations. Ideally, a complete set of biocatalytic rules would be able to spawn all possible enzyme mechanisms, when chained together.

Figure 1 summarises the process we followed for the creation of the rules. First, we parsed the 2D curly arrow diagrams of all the steps and mechanisms in the database to extract the relevant information about bond changes and intervening chemical groups (figure 1-A). Second, a graph was built representing possible electron pathways (figure 1-B) based on the curly arrows and the atoms they touch (up to two bonds away). Finally, by traversing this graph, we extract possible rules that are based on the literature knowledge (figure 1-C). This process leads to the creation of two types of rules: Single-step rules, which are observed in their entirety in at least one mechanistic step in the database; and mixed-step rules, which cannot be found in the database, but are rather the result of combining information from different steps, which share similar chemical groups (see methods). The total number of biocatalytic rules obtained by following this procedure is 7218. Of these, 3676 are single-step rules, while the remaining 3542 arise from mixing information from more than one step (mixed-step rules).

The number of biocatalytic rules, as well as their specificity, is dependent on the exact algorithm followed during their creation. Before settling on the current definition, we tested other ways of building the rules, such as including only the reaction centres (atoms directly involved in bond formation and cleavage) and the immediate surrounding atoms, which leads to the creation of fewer rules. Smaller and less specific rules like these, match more enzyme active sites but lead to spurious matches. For example, these rules are not able to differentiate between a proton transfer from a hydroxyl group and a carboxylic group. On the other hand, rules that include three or more shells around the reaction centres produce more meaningful matches but, by being overly specific, they match fewer active sites and are less useful for searching as a result. We found that rules that include atoms up to two bonds away from the reaction centre are a good compromise between specificity and applicability. We further refined this definition by considering that rules should not discriminate between carbon or hydrogen atoms that are two bonds way from the reaction centres. This means that formic acid and acetic acid, for example, are equivalent for rule matching purposes, when the reaction centre is the negatively charged oxygen. In Figures 1 and 2, positions that can match both C and H are indicated with the “C/H” label.

**Figure 2.**
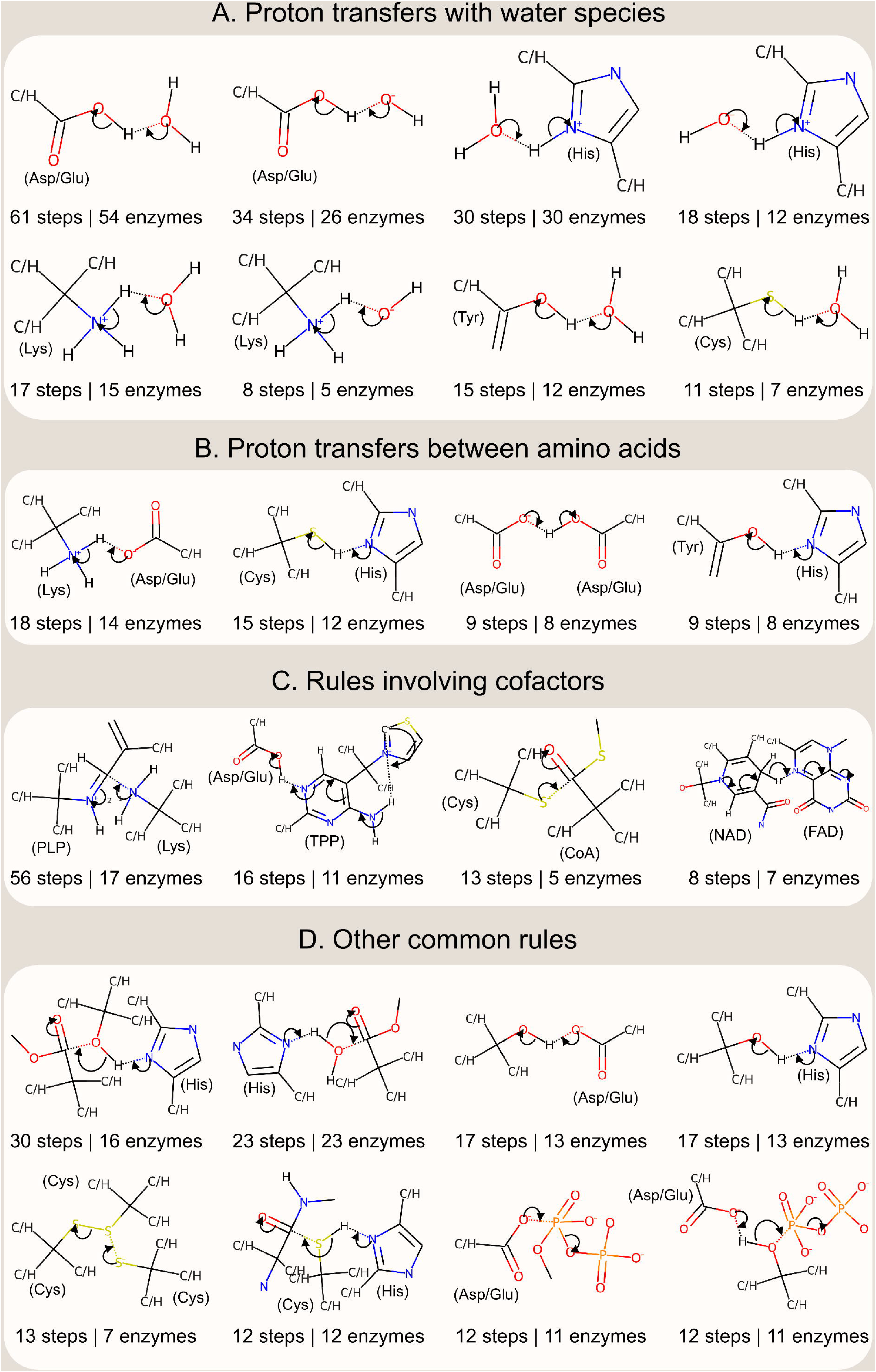
The catalytic rules most commonly observed in the M-CSA database. Rules match chemical groups, not amino-acids or other molecules but, to facilitate comprehension, some molecules and amino acids that match these chemical groups are indicated in parentheses. Glu – Glutamate, Asp – Aspartate, His – Histidine, Lys – Lysine, Tyr – Tyrosine, Cys – Cysteine, PLP - Pyridoxal 5’-phosphate, TPP - Thiamine(1+) diphosphate, CoA - acetyl-coenzyme A, NAD - nicotinamide adenine dinucleotide, FAD - flavin adenine dinucleotide.

The generated rules can be understood as a list of chemical groups that, if present in the active site, can be transformed in a certain way. For example, the first rule shown in Fig 2-B means that if there is a protonated amine group and a deprotonated carboxylic group in the active site, then a proton transfer can occur between the two groups. Rules do not contain any metadata about the location of these chemical groups, which means they can match any molecules regardless of their role in the mechanism (such as catalytic residues, substrates, or cofactors), although in this example they will match mostly Lysine and Aspartate/Glutamate residues. Rules are stored in the database as reaction SMARTS expressions plus an additional column with information about the curly arrows representing the movement of electrons, which allows for the reconstruction of the rule in a pictorial form. In this way, rules are both machine readable and interpretable by humans.

Figure 2 shows the most common biocatalytic rules identified in M-CSA. Although most rules have a corresponding reverse rule (where the reactants and products are reversed), these are not shown in the picture for simplicity. The full set of rules can be browsed at www.ebi.ac.uk/thornton-srv/m-csa/rules/, which also shows the catalytic steps from which each individual rule was extracted (see figure SI −10). In agreement with our previous analysis on the roles of catalytic residues^6^, the most common catalytic rules are involved with proton transfers. The most common one, observed in 61 mechanistic steps and 54 enzymes throughout the database, is the proton transfer between a carboxylic group and a water/hydronium molecule (Figure 2-A). This is a common reaction step that represents the protonation/deprotonation of Asp and Glu amino acids by bulk water, which is often necessary to recycle the active site. A similar rule corresponds to the proton transfer between a carboxylic group and water/hydroxide. In fact, many common rules involve proton transfers between water molecules (to generate hydroxide/water/hydronium) and chemical groups matching specific amino acids, such as imidazole ring (histidine), methylamine (lysine), propan-2-ol (tyrosine), or thiol group (cysteine). Proton transfers between chemical groups that match pairs of amino acids are also common as can be appreciated in figure 2-B for: Lys and Asp/Glu; Cys and His; Asp/Glu to another Asp/Glu; or Tyr to His. Interestingly, although the third and fourth rules in panel D of figure 2 look like proton transfers between Ser to Asp/Glu and Ser to His, respectively, they come mostly from another molecule’s hydroxyl groups, rather than serine residues.

The second most common rule, present in 56 steps and 18 mechanisms, represents a particularly specific reaction, the attack of an amine group on a Pyridoxal 5’-phosphate cofactor (see figure 2-C). Indeed, cofactors tend to perform the same function across enzymes^20^, so other common rules contain groups found in cofactors, such as Thiamine diphosphate and Coenzyme A. However, it is not always the case that the same cofactor will follow the same catalytic rule in different enzymes. NAD(P), for example, is a common cofactor, present in more than 30 steps, but since it donates/receives a hydride from different chemical groups, it is present in many different rules. The most common one, seen in 8 steps, is a hydride transfer between NAD(P) and FAD.

The rules discussed up to now are all single-step rules. Mixed-step rules expand the applicability of EzMechanism by mixing information from different mechanistic steps. The rationale for this is that while interacting curly arrows (representing close chemical groups or molecules) are specific to each other, this might not be the case for chemical groups or molecules that are far away in a curly arrow chain. The bottom part of figure 1-C shows four examples of extended rules with indication of which curly arrows come from the two different steps from which they were extracted.

We hope it becomes apparent from the present discussion that biocatalytic rules, as formulated here, are a powerful tool to understand how enzymes operate. Furthermore, the modular approach we took for decomposing and assembling the rules mimics the evolutionary process in enzymes, in that mutations of single residues will affect only parts of the curly arrow chains^21^. A more detailed analysis of these rules and their usefulness to understand enzyme evolution is in progress.

### Automatic Proposal of Enzyme Mechanisms

To search for the mechanism of specific enzymes, EzMechanism takes as input a three-dimensional structure of the active site, the overall enzyme reaction, and two 2D schemes of the active site representing the initial (reactants) and final (products) states of the mechanism. The 3D model of the active site can be built from an adequate PDB structure and should include all the catalytic residues (identified by conservation, for example, but when in doubt, more residues can be added) as well as the reactants and cofactors of the reaction (similar analogues of these molecules can be used instead). The 2D schemes are mostly used for visualisation of the search results, but also for the definition of the initial protonation states. A webpage was created to facilitate the creation of these inputs (see figures SI-1 to SI-4).

The software’s search algorithm uses these inputs and the catalytic rules described above to generate a landscape of possible new configurations around the reactants and products configurations of the active site and to ultimately find one or more catalytic paths that connect these two endpoints (details in the online methods). Figure 3 depicts this overall process. Starting with the reactants configuration, every catalytic rule is checked against it, and for each matching rule a new configuration is created (see figure 3-A). The new configurations represent possible first intermediates of the mechanism. The relationship between the starting and newly generated configurations can be conveniently represented as a graph where each configuration (mechanism intermediate) is represented as a node and rule matches (mechanistic steps) are represented as edges. The rounded square in figure 3-B contains the same configurations as panel A but in this schematic format. The number 1 on top of the reactants configuration (orange circle) means that this was the first configuration matched against the rules.

**Figure 3.**
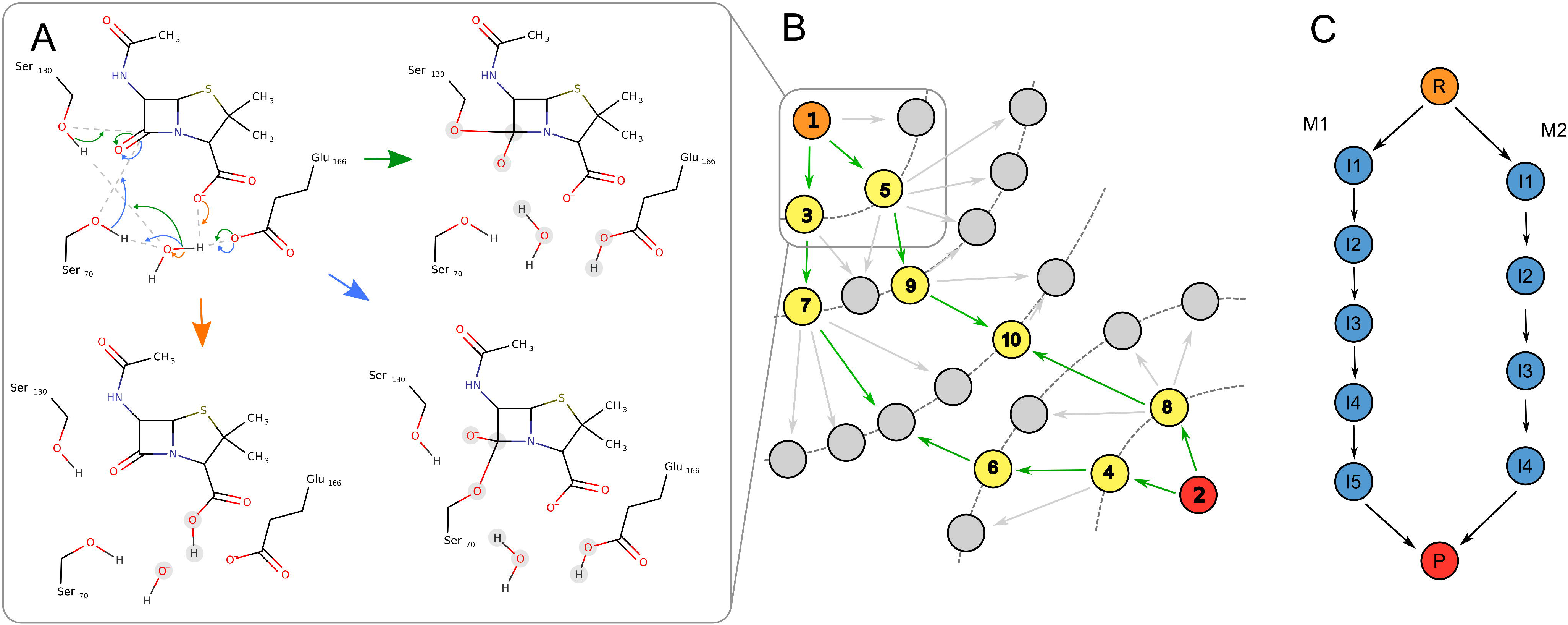
Illustration of the mechanism search algorithm used by EzMechanism. A – Example of a configuration that matches three rules (identified by the three different colours). Each rule match leads to the creation of a new configuration. B – Example of a configuration graph that represents the chemical catalytic space explored by EzMechanism. Green arrow edges represent steps in the search that are prioritised according to the prioritisation algorithm. Numbers in configuration exemplify the order followed by the software to match the rules to each configuration. Orange – Reactant configuration, Red – Products configuration, Yellow – configuration matched with rules, Grey – other configurations. C – A cleaner representation of the two possible mechanisms that are present in B. Every path that links the reactants with the products configuration represents a potential mechanism.

The process of rule matching is similarly performed for the products configuration (red circle in figure 3-B, annotated with number 2) and then in an iterative way for the newly generated configurations (annotated in yellow and with a number denoting the search sequence). Since the number of possible reactions for a typical active site is too big to be explored exhaustively, the search is prioritised towards the most promising configurations (details in the online methods). Grey circles represent configurations that are not checked against the rules. The grey circle inside the rounded square, for example, was generated from a rule matching the reactants configuration, but since that step was unfavourable (indicated by the grey arrow), that part of the chemical space was not explored further.

After matching the rules for a series of configurations, a graph like the example in figure 3-B is generated. In a graph like this, paths of configurations (i.e., sequences of steps) that link reactants with products are possible reaction mechanisms for the enzyme. For this particular example, two potential mechanisms for this enzyme can be identified, which are represented schematically in figure 3-C.

### Validation against known Enzyme Mechanisms

In order to test the applicability of EzMechanism and the quality of its results we started by trying to build a model of the active site of the first 100 entries of the M-CSA. We used a dataset^22^ previously built in our group to quickly identify, among PDB structures of the same enzyme (same UniProt ID), which structures contained ligands that are similar or identical to the cognate ones (the cognate or native ligand of an enzyme is its biological substrate). For almost a third of the initial dataset, 35 entries, we could not build a 3D model of the active site using any PDB structures, due to missing catalytic residues, substrates, or cofactors. This number reflects the low number of structures in the PDB with native ligands and is not a limitation of our software (Only 26% of enzyme structures in the PDB have a ligand with at least 70% similarity to the cognate ligand^22^). EzMechanism could still work for these enzymes but that would require additional modelling to build a suitable enzyme-ligand complex. Another 9 entries were excluded from the analysis because they involved either a radical mechanism (8 enzymes), or a stereoisomerase reaction (1 enzyme), which are out of scope for the current version of the software.

Since the output of the mechanistic search calculation can lead to a large number of configurations with complex associated data, we built a custom webpage to facilitate the analysis of these results (available at www.ebi.ac.uk/thornton-srv/m-csa/predictions/). A table with the overall results of the validation is also included in the SI (table SI-I). In the output page of each prediction, configurations can be manually selected and inspected, and the graph of configurations can be filtered to hide paths larger than a given cut-off or involving bond formation between distant atoms (details in the online methods). The results of the validation for the dataset of 56 enzymes are summarised below. We divided the validation in three sections that test different parts of the program.

We first checked if the algorithm could find the correct mechanism (defined here as the mechanism described in the database) for a given enzyme when using only rules generated from that database entry. Since rules are not extracted directly from the mechanistic steps but are first decomposed into individual curly arrows and later reconstructed (in order to combine information from different steps/enzymes), this test makes sure this process is correctly implemented. It is also a test of the rule matching algorithm as well as the overall mechanistic search. EzMechanism passes this test for 55 of 56 enzymes, failing in an unusual case where there are two independent rules (M-CSA:90 step 2) that share the same atom in the same catalytic step.

The second question we addressed is whether EzMechanism could find the correct mechanism of the remaining 55 enzymes when using the complete rule dataset. Since the rule dataset was created using knowledge from the mechanisms of all enzymes in the M-CSA, including these, this might seem a trivial task. The difficulty arises in the large number of configurations that can be generated for a single active site, which prevents the possibility of doing an exhaustive search across the entire catalytic space. By limiting the number of configurations that are checked against the rules to a maximum of 1000 (yellow circles in figure 3-B), we keep the number of configurations in check, but even so, the number of generated configurations can be larger than 100 000 for some enzymes. This test is then a way of checking how well the prioritisation algorithm works, and how the software handles these large numbers of configurations. EzMechanism is able to find the correct mechanism for 51 enzymes out of the 55 using the default number of explored configurations, and the remaining four mechanisms (which are challenging due to their number of catalytic steps: 8, 9, 10 and 11), can still be found by increasing the number of explored configurations to 10,000.

The final test aims to evaluate the coverage of the catalytic rules. Ideally, the catalytic rules generated from the current knowledge contained in M-CSA would be able to cover not only the enzymes in the database but also uncharacterised enzymes (which nevertheless will share some of the catalytic residues, cofactors, or chemical groups of the substrates). One way to do this analysis is to check how many mechanisms contain “unique” rules i.e., rules that are only observed in that enzyme. If a mechanism contains any step with a unique rule, this means that we could not have guessed this exact mechanism if the enzyme was not already in the database, since these unique rules would not be in the ruleset. 42 of the 55 enzymes contain unique rules. A more relaxed test is checking if a mechanism can still be found for the tested enzymes if the unique rules are not used. In this case, EzMechanism can find a mechanism for 26 of the 55 enzymes under study (in 13 of these cases the mechanism will be different that the one in the database which would require the unique rules). The relevance of these findings is considered in the discussion.

### Test Case: β-Lactamase A

β-Lactamases are bacterial enzymes that hydrolyse the β-Lactam ring present in penicillin and similar antibiotics, conferring resistance to these molecules. Here, we use a class A β-lactamase to exemplify how EzMechanism can be used to explore possible mechanisms.

We prepared a run of EzMechanism based on the 1TEM PDB structure^23^ of the *E. coli* β-lactamase, where a substrate analogue is covalently linked to Ser70, suggesting a nucleophilic role for this residue in the mechanism. Indeed, the accepted reaction mechanism, as described in the M-CSA entry for this enzyme (M-CSA ID:2) involves the creation of this covalent intermediate with the help of Glu166 and a bridging water^24^. This is followed by the cleavage of the four membered lactam ring and a nucleophilic attack by a water molecule, which ultimately leads to the collapse of the covalent intermediate. This mechanism is identified by the green arrows in figure 4.

**Figure 4.**
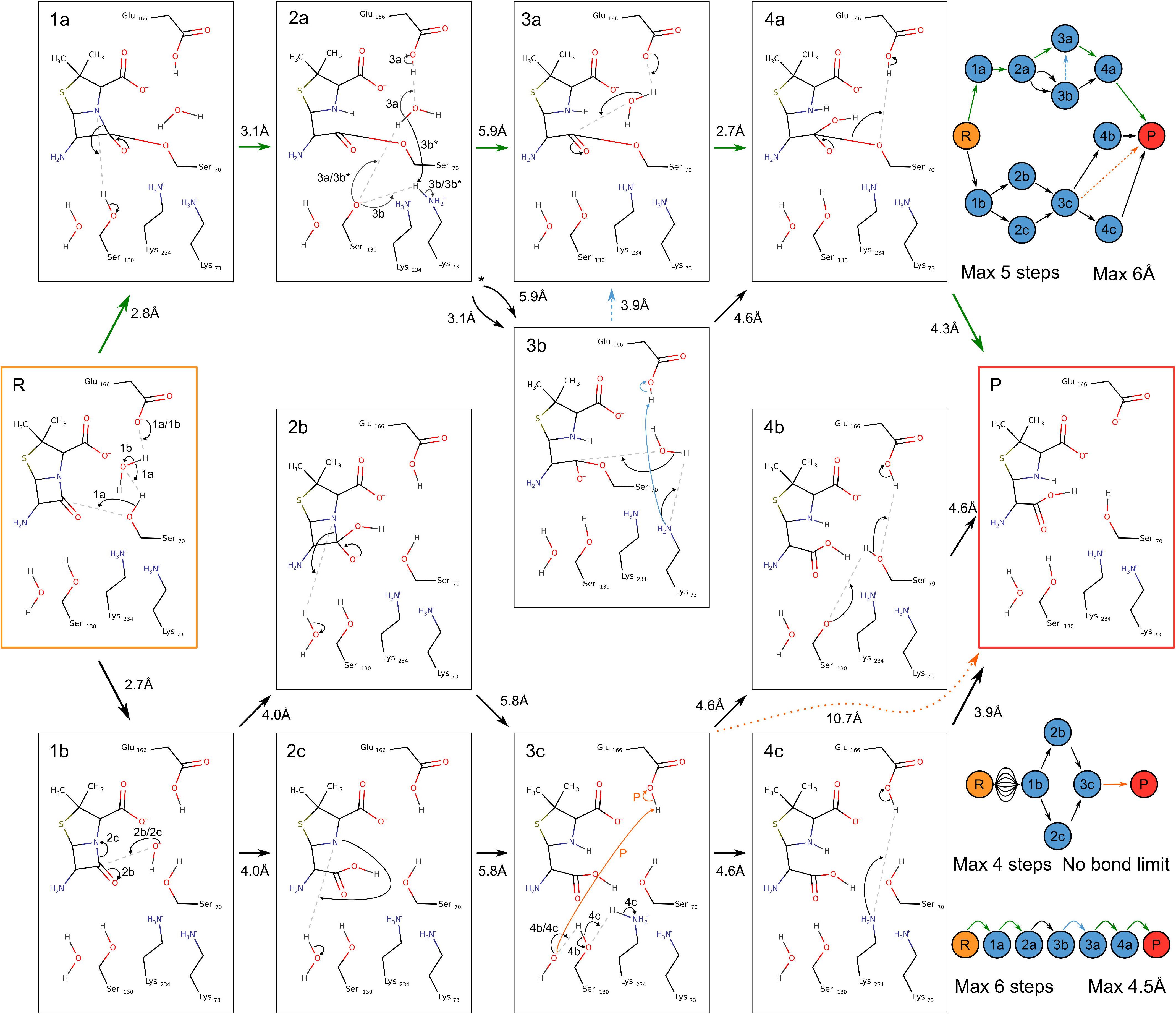
Mechanistic paths found by EzMechanism for β-Lactamase A. Green arrows indicate the mechanism described in the database. The graph of configurations drawn with the 2D curly arrow schemes corresponds to the graph shown on the top right. The two graphs in the bottom right represent other possible paths for the same prediction output that can be generated by considering different cut-offs for the number of steps in the path and the maximum distance of the atoms involved in the formation of new bonds.

The calculation on this active site was limited to 1000 explored configurations (configurations checked against the rules) and generated a total of 9855 configurations. This result is typical for other enzymes we tested, which highlights the complexity and vastness of the chemical space available to enzymes. Although all these configurations are theoretically possible, most of them will not be visited by the enzyme, because they are not energetically favourable. There are computational methods able to calculate the relative energy of the enzyme states along a reaction path, but they are still too computationally expensive to be used in such a large scale. In lieu of the energy, and to facilitate the interpretation of these results, the output page of EzMechanism can use the length of the newly formed bonds (considering the positions of atoms in the PDB structure) and the number of steps in the path to filter less relevant regions of the chemical space.

Figure 4 shows the configurations and paths linking reactants to products that involve a maximum of 5 steps and distances shorter than 6 Å (graph in upper right corner. For simplicity we are hiding conformations steps that involve the carboxylic group of the substrate). The mechanism described for this enzyme in the database (highlighted in green) is one of the possibilities within these criteria. Interestingly, there is an almost identical path, which only differs in two steps and one configuration (2a -> 3b -> 4a, instead of 2a -> 3a -> 4a). In the alternative case, Ser130 is reprotonated by Lys73 (either directly or through a water molecule), and then Lys73 is the nucleophilic base in the following step, rather than Glu166. Although the first of these steps involves closer atoms when using Lys73 (2a -> 3b; 3.1 Å) rather than Glu166 (2a -> 3a; 5.9 Å), this is not the case for the second step (3b -> 4a; 4.6 Å vs 3a -> 4a; 2.7 Å). After investigating the graph of configurations further, with different filters, we found that an additional step transforming 3b into 3a (proton transfer between Lys73 and Glu166) might present a better mechanism alternative, in terms of the distances between new bonds. This alternative mechanism has 6 steps and is shown in the bottom left corner of figure 4.

Finally, there are a couple of alternative paths that start with the deprotonation of a water molecule (R -> 1b), including one possible path with only 4 steps involving a step with two very distant species, represented by the orange arrow). This 4-step path cannot be completely excluded based on the distance criterion, because it involves one hydroxide molecule, which might move in the active site, or be substituted by other water molecules. As for other paths starting with configuration 1b and involving hydroxide, their feasibility can only be ascertained for sure with other methods, such as energy calculations. In other b-lactamase classes with mechanisms involving hydroxide molecules, these are typically stabilised by metal ions.

## Discussion

EzMechanism is a knowledge-based approach to study enzyme mechanisms. It is the first method able to generate mechanistic hypotheses for a given 3D active site in an automated way. EzMechanism has some important advantages when compared with other ways of producing mechanism proposals (which consists mostly of literature searches and human expertise). First, the program ruleset is derived from the mechanisms of hundreds of enzymes belonging to different EC classes and structural superfamilies, which ensures good coverage of the chemical space and surpasses what most humans can recall. Second, because rule matching is purely based on local chemistry at the step level, the program does not limit the search to similar (evolutionary related) enzymes, or enzymes that catalyse the same overall reaction. Thirdly, EzMechanism makes sure that combinations of rules are searched almost exhaustively (guided by the prioritisation algorithm), which might alert the user to paths not previously considered. Finally, all rules and generated catalytic steps link back to the M-CSA entries and the original literature that were used to create them. This facilitates the comparison of the studied mechanisms with the available literature, and the integration of the new studies with existing knowledge.

EzMechanism is not yet a complete solution to predicting enzyme mechanisms *ab-initio*, however. While the search and prioritisation algorithms are able to find the correct mechanisms even when the chemical space is quite large, they are limited by the coverage of the catalytic rules. When excluding from the prediction rules exclusively seen in the enzyme under study, the software can only find a mechanism in about 50% of the cases, and the correct mechanism (defined as the one visible in the database) in about 25%. These should be the expected success rates of EzMechanism to find a complete mechanism, when applied to enzymes not covered by M-CSA. Currently, M-CSA covers 65% of all EC subsubclasses (the third level of the EC classification) and 84% of the subsubclasses with a PDB structure (enzymes that share the same subsubclass typically follow the same mechanism and have a similar substrate). Even for enzymes for which EzMechanism cannot find a complete mechanism, it may still find some of the catalytic steps around the reactants or products configurations, together with the information about the original literature and enzymes that catalyse similar steps.

In future versions of the software, we aim to improve the rule set coverage by adding more mechanisms to the database, include radical reactions to the dataset, and tweak how the chemical information is codified in the reaction smarts patterns to make the rules more generic. Two additional ways to increase the number of rules might involve the manual identification and incorporation of basic rules of organic chemistry, as well as the addition of datasets commonly used for the prediction of reactions in organic syntheses^25^. At the same time, since more general rules will lead to more matches (some of them potentially spurious), we will be working on coupling energy calculations (using QM/MM models) with the predictions, for a more efficient search of the chemical space, and the possibility to assess which reaction paths are more energetically favourable.

Our hope is that, with time, the discussed improvements will allow EzMechanism to completely automate the study of enzyme mechanisms. This is a hefty goal, but one that becomes more urgent as the number of available structures skyrockets. Just recently, the number of experimentally derived structures available in the PDB^26^ surpassed the 200,000 and Alphafold^15^ has released over 200 million predicted structures^16^.

By facilitating the study of enzyme mechanisms, EzMechanism might be useful to other studies and applications where this knowledge is important. For example, it can be used to evaluate the effect of catalytic residue mutations on the mechanism, which should help in the understanding of enzyme evolution, enzyme associated diseases, and the design of new enzyme functions. Additionally, the software can be used to identify potential enzyme reactions when a substrate, native or otherwise, binds an active site. We aim to make EzMechanism available to everyone through a webserver soon, in order to make these and other studies possible.

## Online Methods

### The M-CSA database

The M-CSA (Mechanism and Catalytic Site Atlas) is a manually curated database of catalytic sites and enzyme mechanisms that is freely available at www.ebi.ac.uk/thornton-srv/m-csa/. Currently, M-CSA contains detailed annotations on the enzyme reaction mechanisms of 734 enzymes, including the curly arrow diagrams of 3238 catalytic steps, which are the foundation for the catalytic rules presented in this paper. The data in M-CSA is unique in its scope and breadth, and it has been used by ourselves and others to understand enzyme function and evolution^27–30^. Notably, we would like to highlight the work of Jakob Anderson and colleagues^14^, which used M-CSA data to test a new program they created that is able to perform multi-step chemical reactions *in silico* using graph transformation. For this, and independently from the work described in the present paper, they built a set of catalytic rules that are conceptually similar to our own “single-step” rules. Lastly, they used their software and created rules to find hypothetical mechanisms (i.e. without being mapped to a particular three-dimensional active site) for reactions in RHEA^31^, a database of chemical reactions.

### Programming Details

The code written for this project is integrated with the M-CSA codebase and uses most of the same technologies as the website, which is implemented in the python Django Web Framework (www.djangoproject.com) and uses a PostgreSQL database (www.postgresql.org). The python programming language was used for the code that extracts the chemical rules from the curly arrow diagrams and the search algorithm that tests these rules against the active site configurations. This code makes extensive use of the RDKit Cheminformatics package (www.rdkit.org) to, among other things, manipulate molecule objects, convert molecules into SMARTS, and identifying matches between the chemical rules and active site configurations. We used the python library NetworkX^32^ to create and manipulate the graphs used during the creation of rules for merging all the curly arrow data, and during the mechanistic search to store all the generated active site configurations and reaction paths.

The webpages used to submit new searches and analyse the results use the Django template language to generate the final html pages. The ChemAxon MarvinJS plugin v17.15.0 (www.chemaxon.com) is used in the input webpage to draw the 2D scheme of the active site in the reactants and products conformations. The results page uses Cytoscape JS^33^ to show the graph with the computed mechanistic reaction paths, and custom Javascript code was written to filter and control the data presented in this graph.

### Creation of the Rules of Biocatalysis

The catalytic rules created in this work are based on the curly arrow diagrams of the mechanistic steps in M-CSA (figure 1-A). This data, which includes all the pictured atoms and bonds, as well as the curly arrows that indicate the formation and cleavage of bonds, are stored internally as ChemAxon Marvin files (.mrv extension). Marvin files follow a custom XML schema, which we parse using a custom python script, using the ‘lxml’ python package. Molecules in these files are first recreated as rdkit Mol objects and then converted to a SMARTS string. The SMARTS strings for all molecules in the diagram are then saved in the M-CSA database, together with the information about the curly arrows for each step.

In general terms, a catalytic rule can be thought to have a one-to-one correspondence to a catalytic step. This is the case of the examples shown in figure 1, and it is also what happens when we apply these rules in the search algorithm, i.e., each match of a rule will generate a new configuration that corresponds to a new proposed mechanistic step. This might suggest a straightforward way to create catalytic rules, where each step in the database will be used to create a single catalytic rule. We did not follow this route for a couple of reasons: a) there might be independent chemical activities in the same step, such as two proton transfers (not coupled), which might be better described by two catalytic rules; b) rules created in this way might be overly rigid, in particular when there is a long chain of curly arrows. For example, from step A of figure 1, we know that a water molecule can be activated to perform a nucleophilic attack on phosphate by a histidine. From step B, on the other hand, we learn that a water performing a similar nucleophilic attack can also be activated by a lysine. By using information like this and combining information from different steps we were able to generate more general rules.

In order to mix the information of different steps in the database, we start by representing each chain of curly arrows as a directed graph. Each curly arrow is represented by a node and edges indicate that the curly arrows act sequentially on the same atom or bond. The first and last curly arrows in each chain of arrows are labelled as such. Each node (curly arrow) is defined by the atoms or bonds that it touches and surrounding atoms up to two bonds away. Finally, after repeating this process for each step, all the generated graphs are merged by combining nodes (curly arrows) that carry the same information. Figure 2-A shows a combined curly arrow graph for the pictured catalytic steps.

Starting with the combined graph that contains information of all the curly arrows in the database, rules are created by following this iterative process: a) find all the simple paths between “starting” nodes and “ending” nodes, as well as circular paths, that have 10 or less nodes; b) keep all the paths with curly arrows coming from the same step but remove mixed paths (paths with steps coming from different enzymes) with more than 6 curly arrows. This choice of cut-off guarantees that the number of generated mixed-step rules is manageable, and roughly similar to the number of single-step rules; c) for each path, build all the biocatalytic rules that might be generated by the merging of its curly arrows; d) reverse the reactants and products of the generated rules to also obtain the respective reverse rules.

The rules created in this way are saved to the database as reaction SMARTS and each rule is linked back to the enzymes and mechanistic steps that were used to create it.

### Input Preparation and Webserver Pages

The submission process is done through the M-CSA website and it is divided in two steps, the first to add information about the enzyme active site and chemical reaction, and the second to define the particular parameters of the prediction. Several predictions can be performed for the same enzyme.

The first part of the submission uses the same pages that curators use to edit and update the M-CSA website. These pages can be used to define a reference UniProt sequence and PDB structure, and to choose the catalytic residues in this PDB structure. Following that, any required cofactors can be added as well as the substrates and products of the reaction. The page where the reaction is defined checks if the reaction is balanced using ReactionDecoder^34^.

The second part of the input preparation is specific to a prediction and done through a newly created webpage (figures SI-1 to SI-4). It entails the selection of a PDB file (among a filtered list of structures homologous to the reference PDB chosen in the first step) which should have all the reactants and cofactors or close analogues. Then, the correct PDB chain for each catalytic residue must be chosen (some active sites are formed by catalytic residues coming from several chains), followed by the selection of which molecule in the PDB corresponds to each cofactor and reactant as defined previously in the database. This information will be used to create a 2D scheme of the active site. Crucially, protonation states that differ from the default ones, should be defined in the 2D scheme of reactants and products by adding or removing protons from the adequate molecules.

### The Search Algorithm - Setup

EzMechanism uses diverse kinds of information and molecular representations to navigate through the catalytic chemical space:

- Two-dimensional schemes of the active site that are first created for the reactants and products configurations and later for all the configurations generated during the mechanistic search.
- Three-dimensional representation of the active site, which is used to check distances of formed bonds when creating new active site configurations, as taken from the PDB file selected in the input.
- A SMILES string representation of each active site configuration used to define a unique identifier.
- A balanced RXN file with information about the substrates and products of the overall reaction.

Internally, the search algorithm uses a list of RDKit Mol objects (one for each molecule in the active site, including protein amino acids, the substrate, and cofactors) to store the information associated with each configuration. Each Mol object contains two sets of coordinates that correspond to the 2D and 3D representations of the active site, and the molecular topology information necessary to create the SMILES identifier. All heavy atoms in the active site are labelled with a unique identifier (atom map) so that the algorithm can distinguish between identical molecules (such as two identical catalytic residues for example). Hydrogens atoms, on the contrary, are deemed equivalent, so although the protonation state of molecules is considered, the origin of their protons is irrelevant.

The 3D coordinates attributed to the reactants’ configuration are taken from the selected PDB file. Molecules in the internal representation of the active site (including catalytic residues, cofactors, and substrates) are first mapped to the correct PDB residues, according to the user input. PDB residues often differ from the exact molecules in our model, as structures are usually missing hydrogens and may contain analogues, instead of the native substrates, for example. To overcome this problem, the software superimposes the common parts of the cognate ligand onto the PDB ligand and then optimises the 3D coordinates of the remaining atoms of the cognate ligand using the MMFF94 force field, as implemented in RDKit.

Finally, the RXN file containing the reactants and products of the reaction is used to identify what are the reaction centres of the overall enzyme reaction, i.e., atoms that are involved in the cleavage and formation of bonds. After identifying the reaction centres in the RXN file, the program maps the substrates of the RXN file to the substrates of the reactants’ active site configuration created previously. This information will be used in the prioritisation algorithm, which will favour chemical steps that involve the overall reaction centres.

### The Search Algorithm – Search and Optimisations

The starting point for the search of possible mechanisms is the active site configuration of the reactants. Using the RDkit rdChemReactions module, the software checks every catalytic rule against this configuration, and then generates all possible new configurations based on the matches. These new configurations are possible “first intermediates” of the reaction path, and the rules that matched correspond to potential “first steps” of the mechanism. This process of rule matching is repeated for the products configuration, which, in a reverse manner, leads to the generation of potential “last steps” of the mechanism. The process then continues in an iterative way for the newly generated configurations, leading to an increasing number of configurations and potential mechanistic steps, which represent the chemical space of reactions available for this active site. Ideally, at the end of the run, there will be one or more mechanistic paths that connect the reactant configuration with the products configuration.

While exploring the chemical space around the reactants and products, many possible configurations are generated and some of these are similar or identical to each other. These similar configurations arise from three types of situations: a) a given catalytic rule might match two identical catalytic residues or molecules in the active site, such as a rule that protonates a carboxylic group using water, which will match two Asp residues. In this case, two new similar configurations are possible, where each one of the Asps is protonated in turn; b) a single match to the same exact molecule might lead to several new configurations when the matching molecule contains equivalent atoms. A trivial example is a proton transfer from a protonated Lys side chain which generates three new configurations, one for each proton bound to the terminal nitrogen; c) the same or similar configurations might be found by starting from different configurations and following different rules and mechanistic paths, just by chance. In all these situations, the software needs to decide what should be considered a new or already seen configuration. For this purpose, the algorithm considers that all hydrogen atoms are indistinguishable while heavy atoms are unique (by giving them a unique atom map identifier). This rule avoids the proliferation of superfluous active site configurations created by permutations of hydrogen atoms, as in example b), while considering that residues or other molecules are always unique, even if there is more than one type of the molecule in the active site, as in example a).

In a typical run, the number of potential active site configurations that can be generated (which represents the size of the available chemical space) is too big to be explored in an exhaustive way. For this reason, we have developed a prioritisation algorithm that guides the search towards configurations that are more relevant for the mechanistic path. The prioritisation algorithm uses a score that favours: a) steps where new bonds are formed between close atoms (using 3D coordinates) rather than distant atoms; b) steps where new or cleaved bonds involve the reaction centres of the overall enzyme reaction, as taken from the RXN file; c) by virtue of Dijkstra’s algorithm^35^, which tries to find the shortest path between nodes and computes the overall distance as the sum of scores of every edge (step), steps that are closer to one of the starting configurations are also favoured (details in SI).

As described in the first paragraph of this section, the software starts the search from the reactant’s configuration followed by the product’s configuration. In the subsequent iterations, the search alternates between the two sides of the mechanism, so rather than a reactants-to-products direction, the search is done from both sides simultaneously, taking advantage of the reverse rules, with the aim of find overlapping configurations that join the two sides. This bidirectional search is used to make the search significantly more efficient. In a one-directional, the number of configurations (c) to explore grows exponentially with the number of mechanistic steps (s), 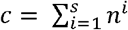, where n is the number of rule matches per configuration (assumed here to be always the same for simplification). With the bidirectional search, however, each side must only explore half of the steps and the formula for the total number of explored configurations becomes 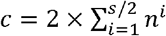. For a six-step mechanism with n=8, for example, the number of generated configurations goes down from roughly 300 000 to around 1000, when comparing the two approaches.

In initial versions of the software, we noticed that most of the computational resources (both CPU and memory) were being spent on creating and saving new configurations and checking the rules against the configurations. We have implemented the following changes to limit the cost of these operations: a) instead of representing each active site configuration as a single object, they are now represented as a list of normalised molecules, meaning that the representation of equivalent molecules in computer memory is shared among configurations; b) the normalisation of molecules across configurations allows for the caching of the rule-matching calculations; c) the rule matching function of RDKit requires the creation of a list of molecules with the same length as the number of molecules in the rule. Instead of just creating all permutations of a given length with all the molecules in the active site, the software first runs a substructure match between each molecule and part of the rule, and the permutations are just created for the matching molecules. The results of the substructure matching are also cached for efficiency.

### Webpage for the analysis of the output

The output of the mechanism search calculations may contain an overwhelming number of active site configurations and reaction steps. To facilitate the interpretation of these results we developed a custom webpage, available at: www.ebi.ac.uk/thornton-srv/m-csa/predictions/ which links to an output page for each of the predictions. The output page contains three main panels, one showing the graph of configurations and steps, another with buttons used to filter the graph and show information about the catalytic rule of the selected reaction step, and a third showing the 2D diagrams of selected configurations (see figures SI-5 and SI-6). If the mechanism prediction is done for a database entry that already contains a mechanism, the mechanism already in the database is shown in a fourth panel for comparison purposes (see figures SI-7, SI-8, and SI-9. We have included a detailed description of the output page and its capabilities in the SI.

