## Supporting Information for "EzMechanism: An Automated Tool to Propose Catalytic Mechanisms of Enzyme Reactions"

### Formula used by the prioritisation algorithm

The formula used to calculate the score of each step (graph edge) used by the prioritisation function has two steps, the first uses the largest bond length of all the new bonds created in that step (max_bond_length).

$$score={(\max\left( 4, max\_bond\_length \right)-3)}^{2}$$

With this formula, bonds up to a length of 4 Å yield the same score of 1. Bonds larger than that are penalised according to a quadratic function.

The second part of the prioritisation is a simple halving of the score for steps whose reaction centres overlap with the reaction centres of the overall reaction. This is a heuristic measure to favour parts of the catalytic space that are moving the reaction forward.

$$score= \frac{score}{2}$$

When choosing the next configuration to match the rules, these scores are used in conjunction with Dijkstra's algorithm, which finds the configuration with the shortest path (using the sum of the scores of each edge as the distance) to the reactant or product configuration. In this way, paths with lower scores are favoured, as well as paths with smaller number of steps.

### Validation of mechanism prediction - Table

Table extracted from: [www.ebi.ac.uk/thornton-srv/m-csa/predictions/](http://www.ebi.ac.uk/thornton-srv/m-csa/predictions/)

Table SI-I. Overall validation results

| **M-CSA ID** | **Prediction ID** | **PDB_ID** | **Results** | **All Rules Shortest Paths Length \| n \| Depth** | **Own Rules Shortest Paths Length \| n \| Depth** | **Other Rules Shortest Paths Length \| n \| Depth** | **Comments** |
| --- | --- | --- | --- | --- | --- | --- | --- |
| 2 | 1 | 1tem | [See results](https://www.ebi.ac.uk/thornton-srv/m-csa/predictions/2/1/output) | 5 \| 32 \| 27 - 342 | 6 \| 28 \| 17 - 382 | 5 \| 32 \| 27 - 342 |  |
| 7 | 1 | 4aj3 | [See results](https://www.ebi.ac.uk/thornton-srv/m-csa/predictions/7/1/output) | 4 \| 10 \| 11 - 73 | 4 \| 6 \| 11 - 73 | 0 \| 0 \| - |  |
| 8 | 1 | 3c2v | [See results](https://www.ebi.ac.uk/thornton-srv/m-csa/predictions/8/1/output) | 7 \| 312 \| 600 - 615 | 7 \| 4 \| 600 - 615 | 0 \| 0 \| - | Tweaked Model PDB: to put N in the right place (substrate is symmetrical) |
| 9 | 1 | 5t8s | [See results](https://www.ebi.ac.uk/thornton-srv/m-csa/predictions/9/1/output) | 3 \| 2 \| 2 | 3 \| 1 \| 2 | 0 \| 0 \| - | Protonated His. |
| 10 | 1 | 1mka | [See results](https://www.ebi.ac.uk/thornton-srv/m-csa/predictions/10/1/output) | 4 \| 1 \| 21 | 4 \| 1 \| 21 | 6 \| 4 \| 26 | protonated AspB |
| 15 | 1 | 4h0d | [See results](https://www.ebi.ac.uk/thornton-srv/m-csa/predictions/15/1/output) | 3 \| 2 \| 2 | 3 \| 1 \| 2 | 3 \| 2 \| 2 |  |
| 16 | 1 | 4h0d | [See results](https://www.ebi.ac.uk/thornton-srv/m-csa/predictions/16/1/output) | 4 \| 2 \| 5 | 4 \| 1 \| 5 | 9 \| 10 \| 194 - 335 | deprotonated product |
| 17 | 1 | 1a9t | [See results](https://www.ebi.ac.uk/thornton-srv/m-csa/predictions/17/1/output) | 3 \| 1 \| 5 | 3 \| 1 \| 5 | 4 \| 2 \| 67 | same molecule involved in two parts of the rule changed tautomerization state of products |
| 21 | 1 | 1pj2 | [See results](https://www.ebi.ac.uk/thornton-srv/m-csa/predictions/21/1/output) | 4 \| 8 \| 11 - 344 | 5 \| 2 \| 24 - 26 | 4 \| 8 \| 11 - 344 |  |
| 23 | 1 | 1kia | [See results](https://www.ebi.ac.uk/thornton-srv/m-csa/predictions/23/1/output) | 2 \| 1 \| 1 | 2 \| 1 \| 1 | 2 \| 1 \| 1 | tweak: changed protonation state of substrate |
| 24 | 1 | 1nzy | [See results](https://www.ebi.ac.uk/thornton-srv/m-csa/predictions/24/1/output) | 5 \| 2 \| 19 | 5 \| 2 \| 19 | 0 \| 0 \| - | Protonated Asp |
| 26 | 1 | 4i9t | [See results](https://www.ebi.ac.uk/thornton-srv/m-csa/predictions/26/1/output) | 3 \| 20 \| 2 - 145 | 3 \| 4 \| 2 | 3 \| 20 \| 2 - 145 | tweak: protonated one of the histidines |
| 28 | 1 | 1djy | [See results](https://www.ebi.ac.uk/thornton-srv/m-csa/predictions/28/1/output) | 3 \| 17 \| 2 - 562 | 3 \| 3 \| 83 - 562 | 3 \| 14 \| 2 | tweak: put proton in phosphate in products. R groups to C and fix isotope Model not the best, big distances |
| 29 | 1 | 1djp | [See results](https://www.ebi.ac.uk/thornton-srv/m-csa/predictions/29/1/output) | 4 \| 4 \| 57 - 138 | 5 \| 2 \| 127 - 138 | 5 \| 529 \| 126 - 625 | tweak: deprotonated amine and protonated aminoacid to be consistent with M29 in products |
| 31 | 1 | 1lce | [See results](https://www.ebi.ac.uk/thornton-srv/m-csa/predictions/31/1/output) | 6 \| 1 \| 446 | 6 \| 1 \| 446 | 0 \| 0 \| - | Changed protonation states of Cys and Asp. works but very large distances throughout (water) |
| 32 | 1 | 1fro | [See results](https://www.ebi.ac.uk/thornton-srv/m-csa/predictions/32/1/output) | 5 \| 4 \| 28 - 30 | 5 \| 2 \| 28 - 30 | 5 \| 4 \| 28 - 30 |  |
| 35 | 1 | 1phk | [See results](https://www.ebi.ac.uk/thornton-srv/m-csa/predictions/35/1/output) | 3 \| 2 \| 2 | 3 \| 2 \| 2 | 3 \| 2 \| 2 | protonate substrate in products |
| 36 | 1 | 1qq6 | [See results](https://www.ebi.ac.uk/thornton-srv/m-csa/predictions/36/1/output) | 4 \| 16 \| 7 - 9 | 4 \| 2 \| 7 - 9 | 0 \| 0 \| - | protonated one of the waters in the products |
| 38 | 1 | 4du6 | [See results](https://www.ebi.ac.uk/thornton-srv/m-csa/predictions/38/1/output) | 11 \| 132 \| 5127 - 7975 | 11 \| 132 \| 5127 - 7975 | 0 \| 0 \| - | 10 steps protonated two his |
| 40 | 1 | 2paa | [See results](https://www.ebi.ac.uk/thornton-srv/m-csa/predictions/40/1/output) | 2 \| 2 \| 1 | 2 \| 1 \| 1 | 2 \| 2 \| 1 |  |
| 43 | 1 | 2qfp | [See results](https://www.ebi.ac.uk/thornton-srv/m-csa/predictions/43/1/output) | 3 \| 2 \| 2 | 3 \| 1 \| 2 | 4 \| 107 \| 13 - 930 | his not protonated at the end, active site not recycled. phosphate with two protons |
| 44 | 1 | 1ew8 | [See results](https://www.ebi.ac.uk/thornton-srv/m-csa/predictions/44/1/output) | 3 \| 39 \| 2 | 4 \| 2 \| 4 | 3 \| 39 \| 2 | Step 4 is a regeneration step. |
| 48 | 1 | 4rht | [See results](https://www.ebi.ac.uk/thornton-srv/m-csa/predictions/48/1/output) | 3 \| 3 \| 2 | 3 \| 3 \| 2 | 0 \| 0 \| - |  |
| 50 | 1 | 3uwq | [See results](https://www.ebi.ac.uk/thornton-srv/m-csa/predictions/50/1/output) | 4 \| 2 \| 7 | 4 \| 2 \| 7 | 0 \| 0 \| - |  |
| 51 | 1 | 1os1 | [See results](https://www.ebi.ac.uk/thornton-srv/m-csa/predictions/51/1/output) | 3 \| 1 \| 2 | 3 \| 1 \| 2 | 3 \| 1 \| 2 |  |
| 52 | 1 | 3q94 | [See results](https://www.ebi.ac.uk/thornton-srv/m-csa/predictions/52/1/output) | 4 \| 1 \| 4 | 4 \| 1 \| 4 | 4 \| 1 \| 4 | fixed position of the hydroxide in the products conformation of 3q94 not good, Glu too far away |
| 53 | 1 | 2gq3 | [See results](https://www.ebi.ac.uk/thornton-srv/m-csa/predictions/53/1/output) | 4 \| 4 \| 14 - 377 | 4 \| 4 \| 14 - 377 | 0 \| 0 \| - | protonated products |
| 54 | 1 | 3m7w | [See results](https://www.ebi.ac.uk/thornton-srv/m-csa/predictions/54/1/output) | 9 \| 424 \| 374 - 8653 | 10 \| 8 \| 1261 - 6550 | 0 \| 0 \| - |  |
| 55 | 1 | 1gu1 | [See results](https://www.ebi.ac.uk/thornton-srv/m-csa/predictions/55/1/output) | 4 \| 6 \| 7 | 4 \| 1 \| 7 | 0 \| 0 \| - | Fixed protonation of Tyr and Glu |
| 57 | 1 | 3a8o | [See results](https://www.ebi.ac.uk/thornton-srv/m-csa/predictions/57/1/output) | 5 \| 5 \| 28 - 177 | 8 \| 16 \| 65 - 88 | 0 \| 0 \| - |  |
| 58 | 1 | 5o5k | [See results](https://www.ebi.ac.uk/thornton-srv/m-csa/predictions/58/1/output) | 3 \| 4 \| 2 | 3 \| 2 \| 2 | 3 \| 2 \| 2 |  |
| 60 | 1 | 1ne7 | [See results](https://www.ebi.ac.uk/thornton-srv/m-csa/predictions/60/1/output) | 9 \| 18 \| 4063 | 9 \| 18 \| 4063 | 0 \| 0 \| - | 8 steps |
| 61 | 1 | 1bjp | [See results](https://www.ebi.ac.uk/thornton-srv/m-csa/predictions/61/1/output) | 3 \| 1 \| 2 | 3 \| 1 \| 2 | 0 \| 0 \| - | fixed 2d drawing of n-terminal proline |
| 65 | 1 | 5k7x | [See results](https://www.ebi.ac.uk/thornton-srv/m-csa/predictions/65/1/output) | 6 \| 13 \| 31 - 625 | 6 \| 3 \| 31 - 625 | 0 \| 0 \| - | protonated his and deprotonated substrate |
| 67 | 1 | 1pjc | [See results](https://www.ebi.ac.uk/thornton-srv/m-csa/predictions/67/1/output) | 5 \| 2 \| 313 | 5 \| 1 \| 313 | 6 \| 3 \| 160 | changed protonation state substrate |
| 69 | 1 | 2qia | [See results](https://www.ebi.ac.uk/thornton-srv/m-csa/predictions/69/1/output) | 3 \| 1 \| 2 | 3 \| 1 \| 2 | 3 \| 1 \| 2 |  |
| 73 | 1 | 1dcp | [See results](https://www.ebi.ac.uk/thornton-srv/m-csa/predictions/73/1/output) | 5 \| 2 \| 21 - 48 | 5 \| 2 \| 21 - 48 | 0 \| 0 \| - | protonated two histidines |
| 74 | 1 | 1dam | [See results](https://www.ebi.ac.uk/thornton-srv/m-csa/predictions/74/1/output) | 5 \| 2 \| 41 - 137 | 5 \| 2 \| 41 - 137 | 0 \| 0 \| - |  |
| 75 | 1 | 12as | [See results](https://www.ebi.ac.uk/thornton-srv/m-csa/predictions/75/1/output) | 4 \| 5 \| 11 | 4 \| 4 \| 11 | 4 \| 1 \| 11 | protonated ammonia |
| 77 | 1 | 4ubv | [See results](https://www.ebi.ac.uk/thornton-srv/m-csa/predictions/77/1/output) | 5 \| 2 \| 33 | 5 \| 2 \| 33 | 0 \| 0 \| - |  |
| 78 | 1 | 3csc | [See results](https://www.ebi.ac.uk/thornton-srv/m-csa/predictions/78/1/output) | 4 \| 2 \| 9 - 67 | 4 \| 2 \| 9 - 67 | 0 \| 0 \| - | protonated products |
| 79 | 1 | 1jho | [See results](https://www.ebi.ac.uk/thornton-srv/m-csa/predictions/79/1/output) | 2 \| 1 \| 1 | 2 \| 1 \| 1 | 0 \| 0 \| - | fix aromaticity of substrate and which N binds protonated glu in products |
| 80 | 1 | 2x75 | [See results](https://www.ebi.ac.uk/thornton-srv/m-csa/predictions/80/1/output) | 4 \| 8 \| 41 - 222 | 4 \| 1 \| 41 | 0 \| 0 \| - | protonated one of the histidines |
| 81 | 1 | 4csm | [See results](https://www.ebi.ac.uk/thornton-srv/m-csa/predictions/81/1/output) | 2 \| 1 \| 1 | 2 \| 1 \| 1 | 2 \| 1 \| 1 |  |
| 83 | 1 | 1l8s | [See results](https://www.ebi.ac.uk/thornton-srv/m-csa/predictions/83/1/output) | 2 \| 1 \| 1 | 3 \| 1 \| 2 | 2 \| 1 \| 1 | protonated products |
| 84 | 1 | 1b66 | [See results](https://www.ebi.ac.uk/thornton-srv/m-csa/predictions/84/1/output) | 7 \| 8 \| 5634 | 8 \| 4 \| 3992 - 4218 | 0 \| 0 \| - |  |
| 85 | 1 | 1ik4 | [See results](https://www.ebi.ac.uk/thornton-srv/m-csa/predictions/85/1/output) | 5 \| 6 \| 113 - 493 | 5 \| 2 \| 113 | 7 \| 6 \| 91 - 98 |  |
| 86 | 1 | 4bg4 | [See results](https://www.ebi.ac.uk/thornton-srv/m-csa/predictions/86/1/output) | 3 \| 10 \| 2 | 3 \| 2 \| 2 | 0 \| 0 \| - | deprotonated cysteine protonated product |
| 87 | 1 | 5ol4 | [See results](https://www.ebi.ac.uk/thornton-srv/m-csa/predictions/87/1/output) | 5 \| 24 \| 13 - 282 | 5 \| 8 \| 13 - 282 | 0 \| 0 \| - |  |
| 90 | 1 | 5utu | [See results](https://www.ebi.ac.uk/thornton-srv/m-csa/predictions/90/1/output) | 0 \| 0 \| - | 0 \| 0 \| - | 0 \| 0 \| - | protonated asp137 nad to nad+ |
| 91 | 1 | 3h5q | [See results](https://www.ebi.ac.uk/thornton-srv/m-csa/predictions/91/1/output) | 5 \| 5 \| 4 - 21 | 5 \| 1 \| 4 | 7 \| 26 \| 110 - 914 |  |
| 95 | 1 | 1fui | [See results](https://www.ebi.ac.uk/thornton-srv/m-csa/predictions/95/1/output) | 4 \| 23 \| 4 - 381 | 5 \| 2 \| 92 - 184 | 4 \| 21 \| 4 - 23 |  |
| 96 | 1 | 1chm | [See results](https://www.ebi.ac.uk/thornton-srv/m-csa/predictions/96/1/output) | 5 \| 1 \| 103 | 5 \| 1 \| 103 | 0 \| 0 \| - | deprotonated product protonated second h2o |
| 97 | 1 | 4eg2 | [See results](https://www.ebi.ac.uk/thornton-srv/m-csa/predictions/97/1/output) | 5 \| 2 \| 9 | 6 \| 1 \| 26 | 5 \| 2 \| 9 |  |
| 98 | 1 | 5kob | [See results](https://www.ebi.ac.uk/thornton-srv/m-csa/predictions/98/1/output) | 4 \| 3 \| 3 - 12 | 4 \| 1 \| 3 | 0 \| 0 \| - | changed protonation state of two products |
| 100 | 1 | 5jry | [See results](https://www.ebi.ac.uk/thornton-srv/m-csa/predictions/100/1/output) | 5 \| 2 \| 46 | 6 \| 8 \| 28 - 30 | 5 \| 2 \| 46 | deprotonated cys protonated glu protonated water in product (extra proton) |

### Description of the output page

The output page for each of predictions made is available at: www.ebi.ac.uk/thornton-srv/m-csa/predictions/. The output page contains three main panels, one showing the graph of configurations and steps, another with buttons used to filter the graph and show information about the catalytic rule of the selected reaction step, and a third showing the 2D diagrams of selected configurations (see figures SI-5 and SI-6). If the mechanism prediction is done for a database entry that already contains a mechanism, the mechanism already in the database is shown in a fourth panel for comparison purposes (see figures SI-7, SI-8, and SI-9. In the SI, we have included a detailed description of the output page and its capabilities.

On the graph shown on the top left (figure SI-5 and SI-6), configurations are represented as circles and reaction steps, which correspond each to a single catalytic rule, as the edges that connect two configurations. The reactants and products configurations are coloured in orange and red, respectively. Configurations that were matched against the catalytic rules are shown in yellow, and the remaining configurations, which were generated but not checked against the rules are coloured in grey. The configurations are labelled with a unique identifier that starts with a number representing the distance to the reactants’ configuration, followed by a string of letters that hold no particular meaning. Edges are typically coloured in grey. However, if the prediction is for a mechanism already existing in the database, they are coloured in red if the rule is exclusive to that database mechanism and coloured in orange if the rule is seen in this mechanism but also elsewhere.

The two-dimensional scheme of any configuration can be seen below the graph if the corresponding circle is selected. Similarly, if an edge is clicked, two configurations are shown, corresponding to the starting and ending points of that reaction step. In this case, the reaction centres of this step are highlighted in both schemes. The direction of the reaction step can be reversed for visualisation purposes by clicking the edge a second time. In the right side, every time a reaction step is selected, the catalytic rule correspondent to that reaction step is also shown, together with some metrics relative to the prioritisation algorithm and a link to that rule page (see figure SI-10 for an example). When an edge is selected, all the edges for reaction steps that follow the same rule have their representation changed to a dashed line.

The purpose of the buttons and slides on the right side of the output page is to filter and trim the mechanism graph to facilitate comprehension. The three red buttons allow for the removal of any selected configuration, reaction step, or reaction rule (which removes all the reaction steps that follow the selected rule) from the graph. Any elements of the graph deleted in this manner are listed and can be restored. Below the buttons there is a slider to filter out any reaction steps that involve the formation of bonds between atoms farther away than the selected cut-off. Note that the distances are based on the position of the atoms in the PDB structure, so the formation of a bond between atoms that are 7 Å away for example, might not be unreasonable, since during the reaction the molecules in the active site might move to bring these atoms closer. Below the slider, there are four self-explanatory checkboxes to further limit the number of circles or edges in the graph. The second and third (“keep only rules from mechanism”, and “hide rules unique to mechanism”) are only relevant when the prediction is for a database entry with a curated mechanism. Finally, the layout of the graph might be toggled between tree or network-like. The tree layout is particularly useful to see the length of reaction paths, and how far away each configuration is from the reactants. The network-like layout is useful to detect interesting shapes in the topology of the graph, such as bottleneck configurations or reaction steps that are essential to connect the reactants to the products.

If there is a mechanism in the database already annotated for the entry for which the prediction was done, this mechanism is shown below (figures SI-7, SI-8, and SI-9). The 2D curly arrow diagram of every step of the mechanism is shown on the left side of the panel, so it can be compared with the output of the prediction. On the right side of the 2D schemes, the catalytic rules that were originated from each step are shown, and it is indicated if those rules were included in the prediction and matched any configuration during the search.

### Figures

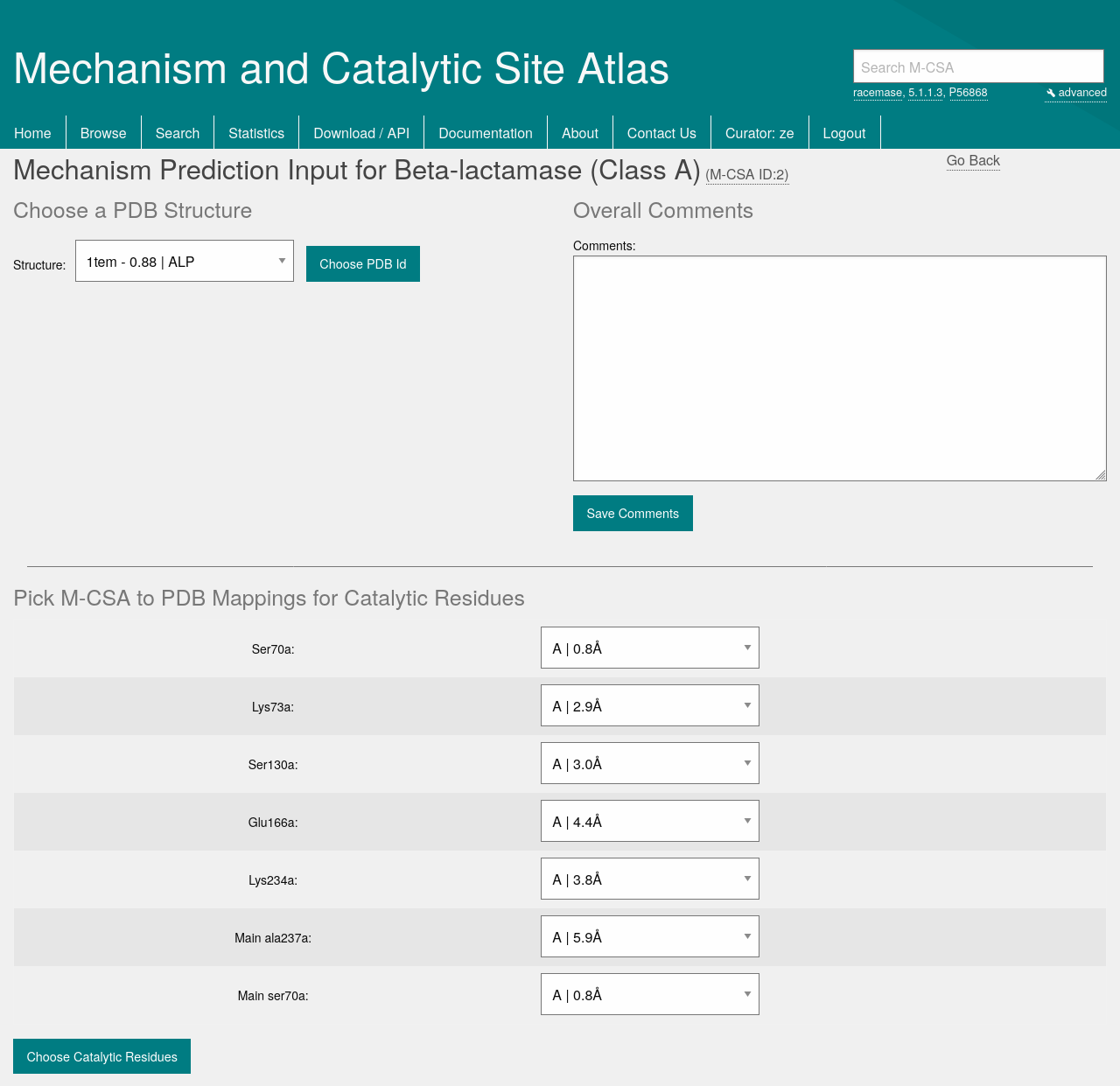

Figure SI 1 – Section of the input page with a form to select a PDB structure of the enzyme under study and another form to define the active site by selecting the appropriate chain of each catalytic residue.

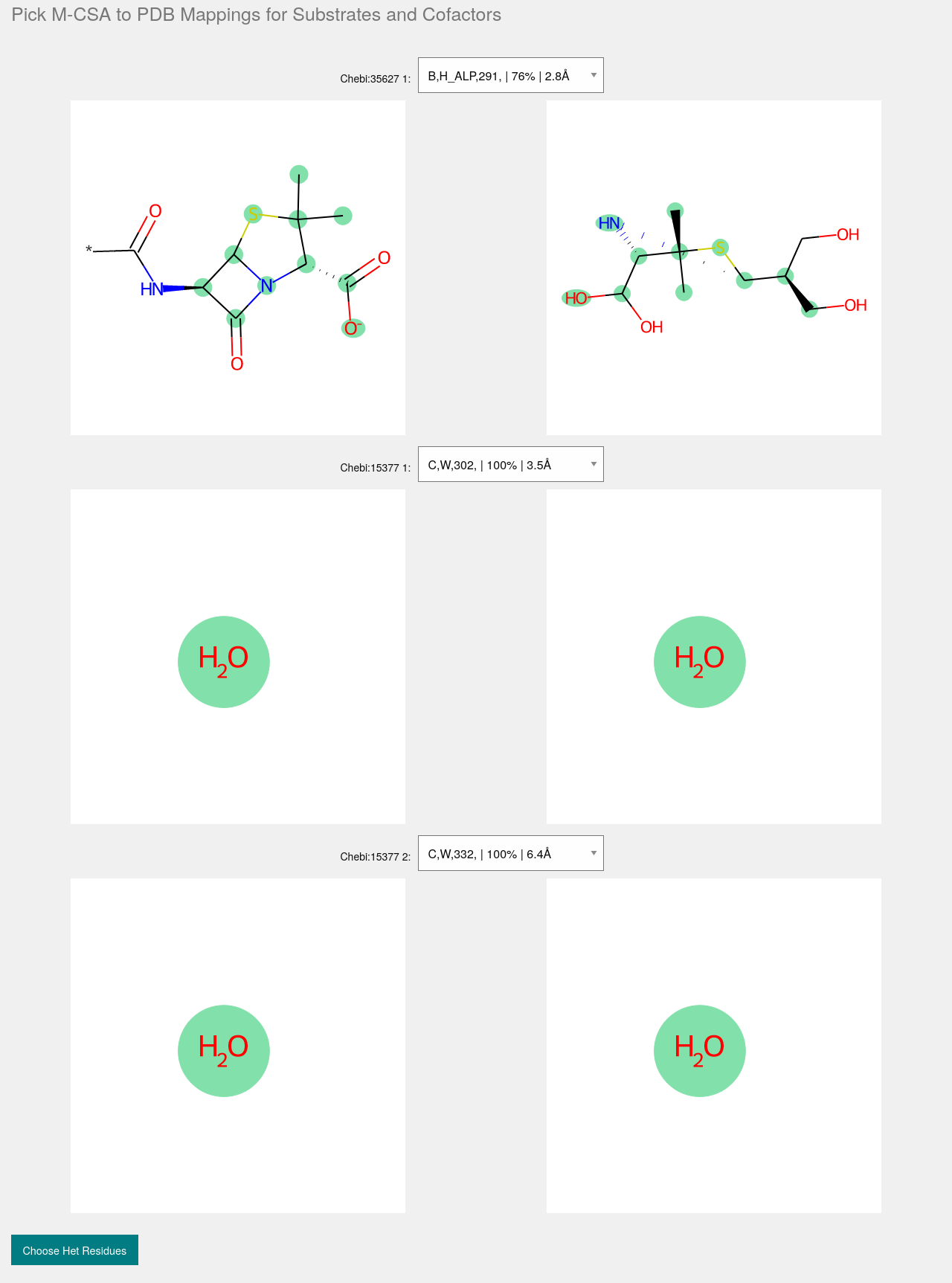

Figure SI 2 – Section of the input page to select the mappings between the native enzyme ligands (substrates and cofactors) and the PDB ligands.

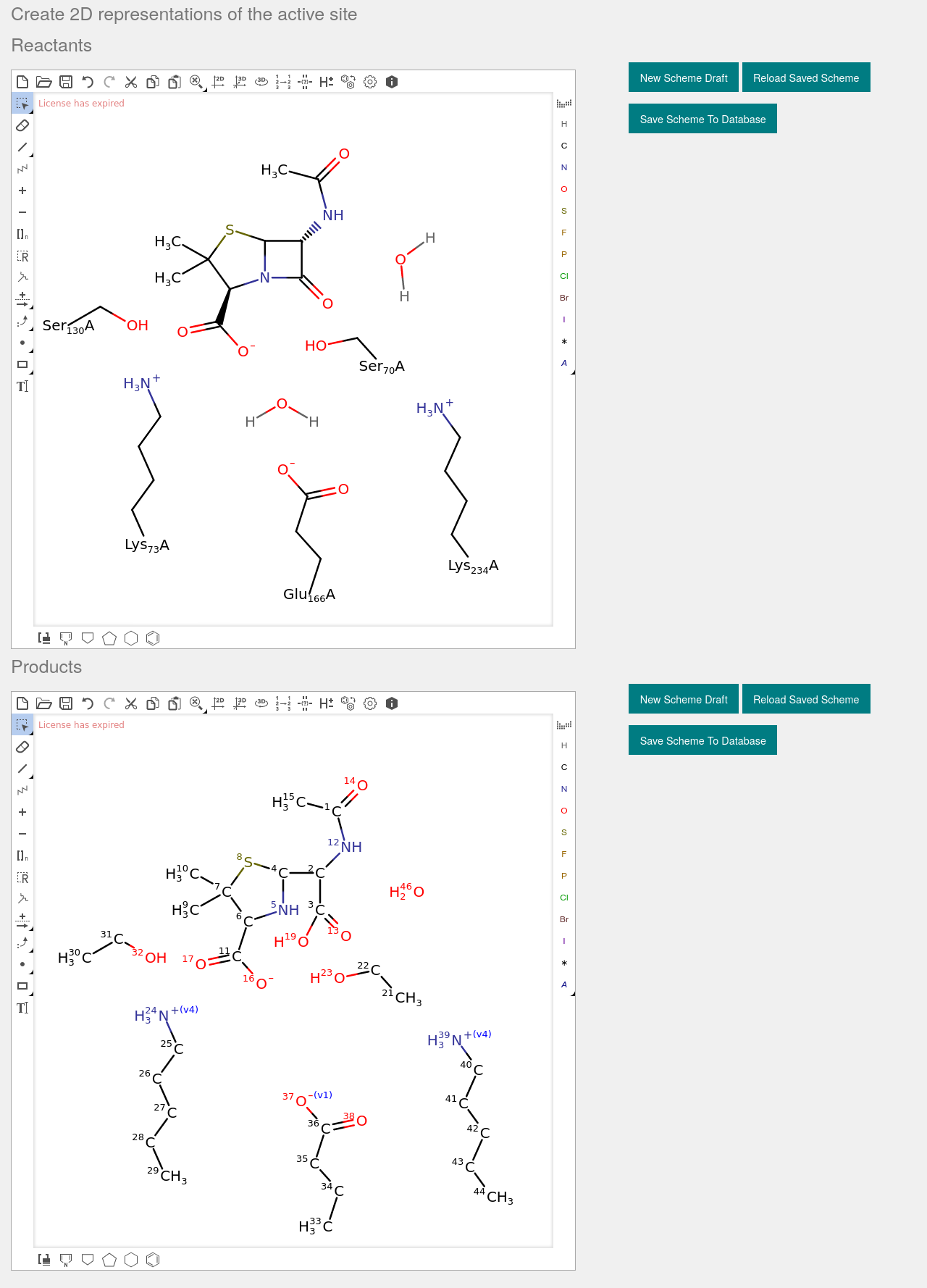

Figure SI 3 – Section of the input page with the MarvinJS plugin where the configurations of reactants and products can be drawn.

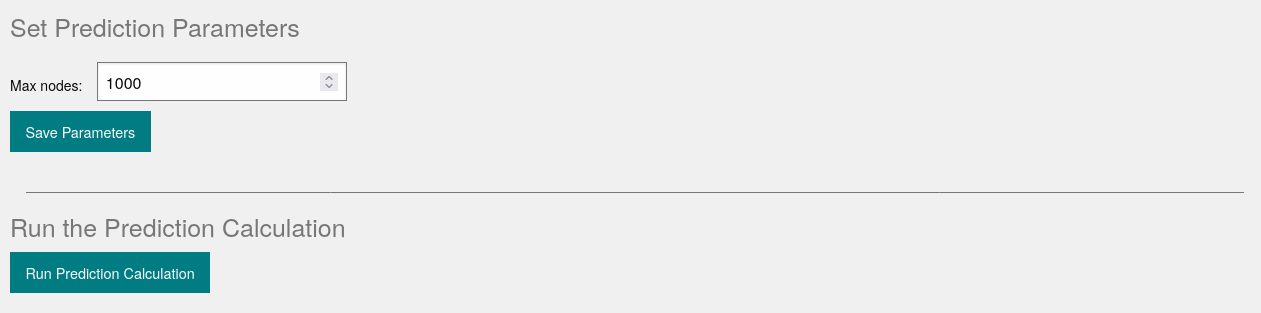

Figure SI 4 – Section of the input page where parameters of the calculation can be edit and the calculation can be submitted.

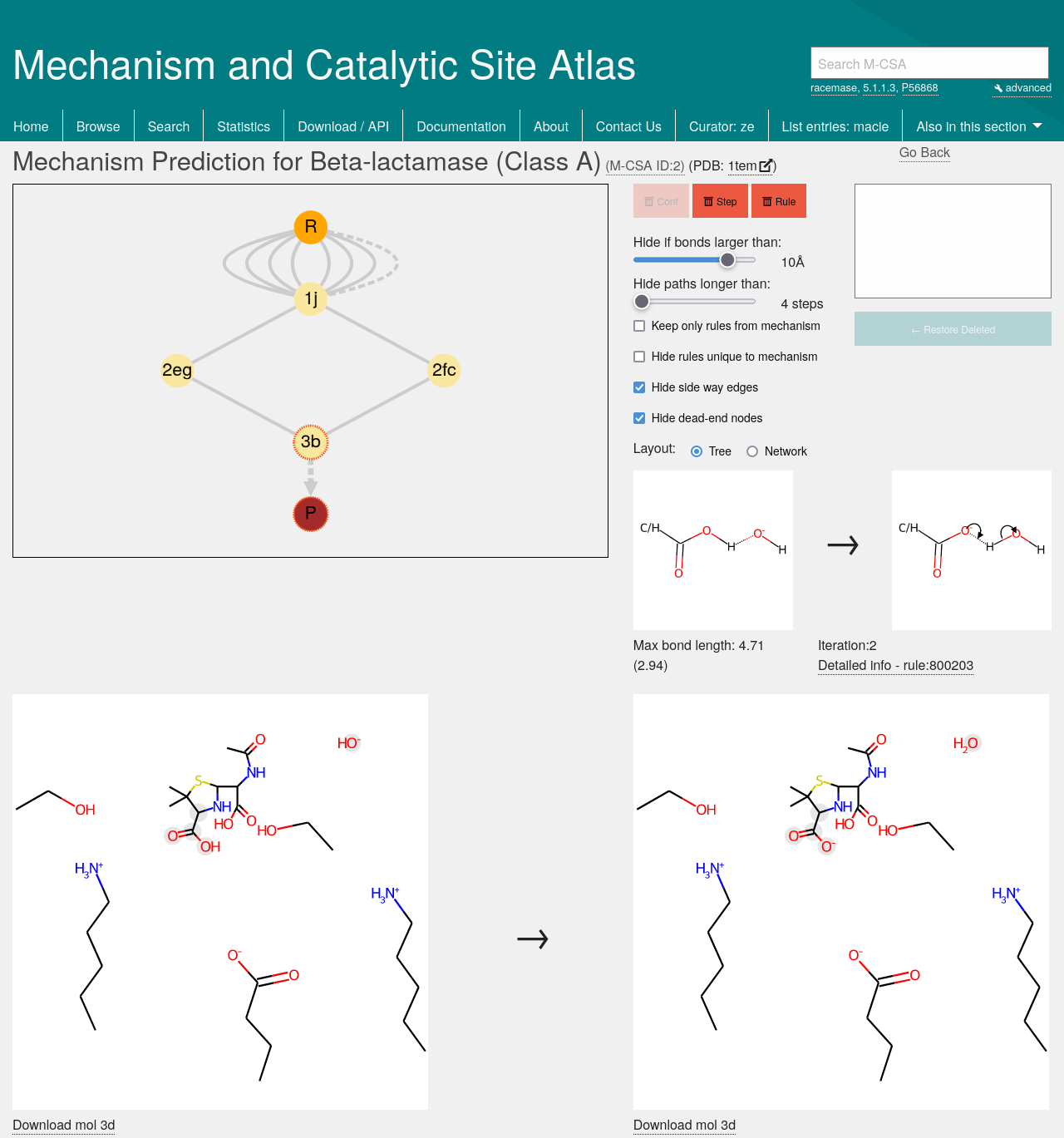

Figure SI 5 – Output page of EzMechanism showing the results of the β-Lactamase A calculation when limiting the length of the mechanism path to 4 steps and the maximum bond distance to 10 Å. The last step of the shown mechanisms (3b -> P) is selected so configurations 3b and P are shown as 2D diagrams on the bottom part of the picture. The rule associated with this step is also shown on the right.

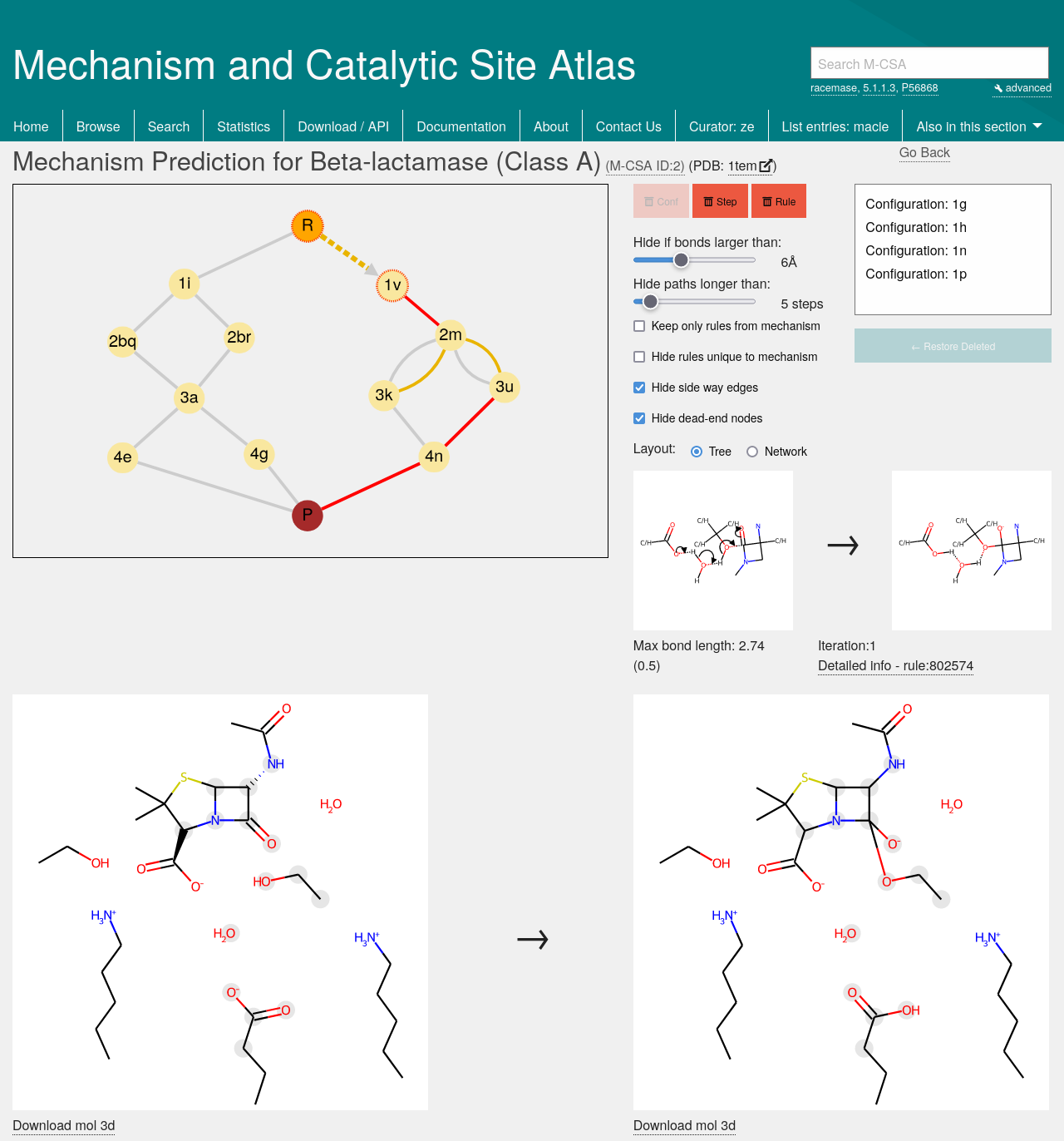

Figure SI 6 - Output page of EzMechanism showing the results of the β-Lactamase A calculation when limiting the length of the mechanism path to 5 steps and the maximum bond distance to 6 Å. The first step of one of the mechanisms (R -> 1v) is selected so configurations R and 1v are shown as 2D diagrams on the bottom part of the picture. The rule associated with this step is also shown on the right.

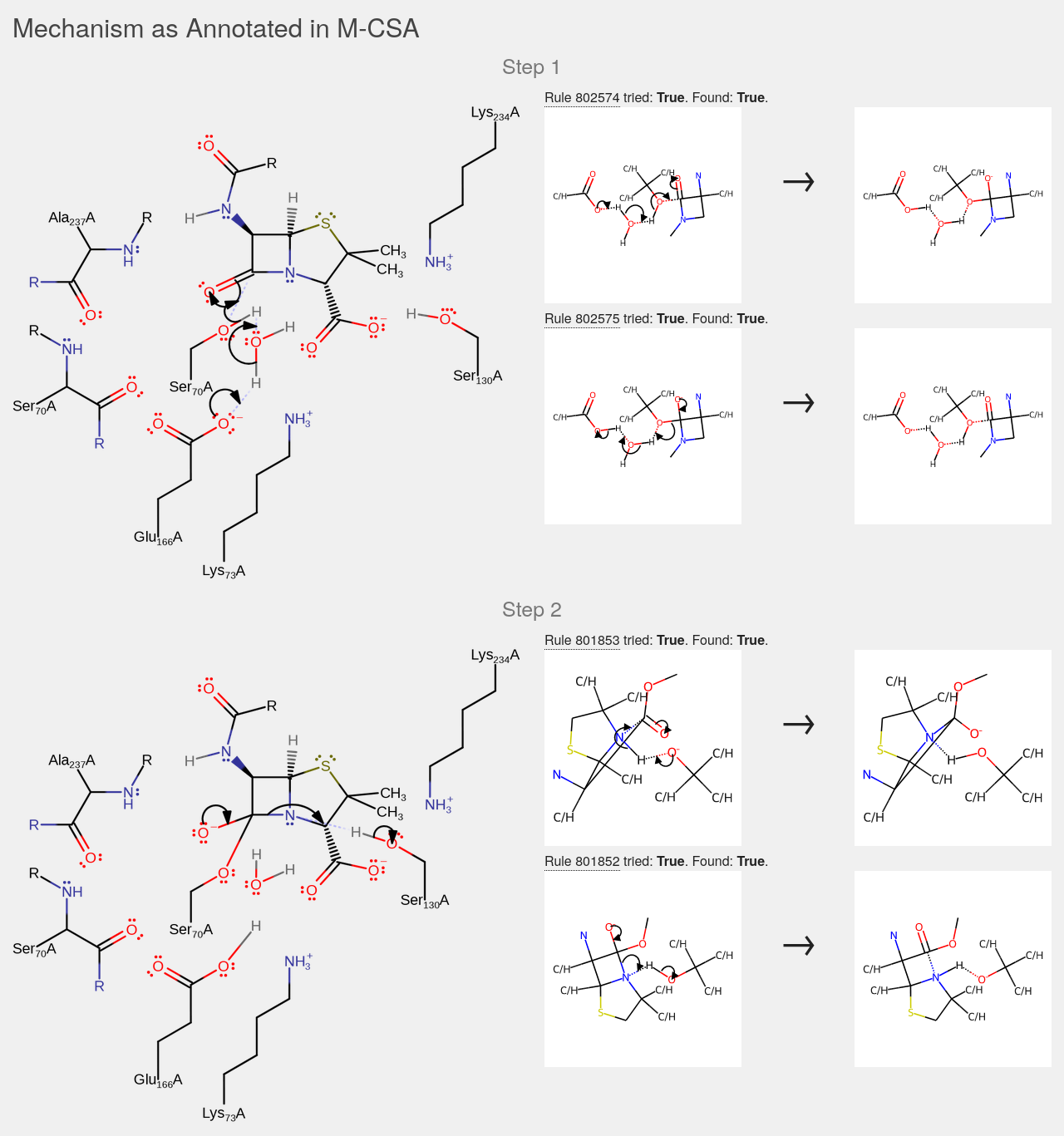

Figure SI 7 – The first two steps of the mechanism of β-Lactamase A as annotated in the M-CSA. Rules extracted from each step are shown on the right as well as an indication of they were used in the prediction and if they were matched to any configuration.

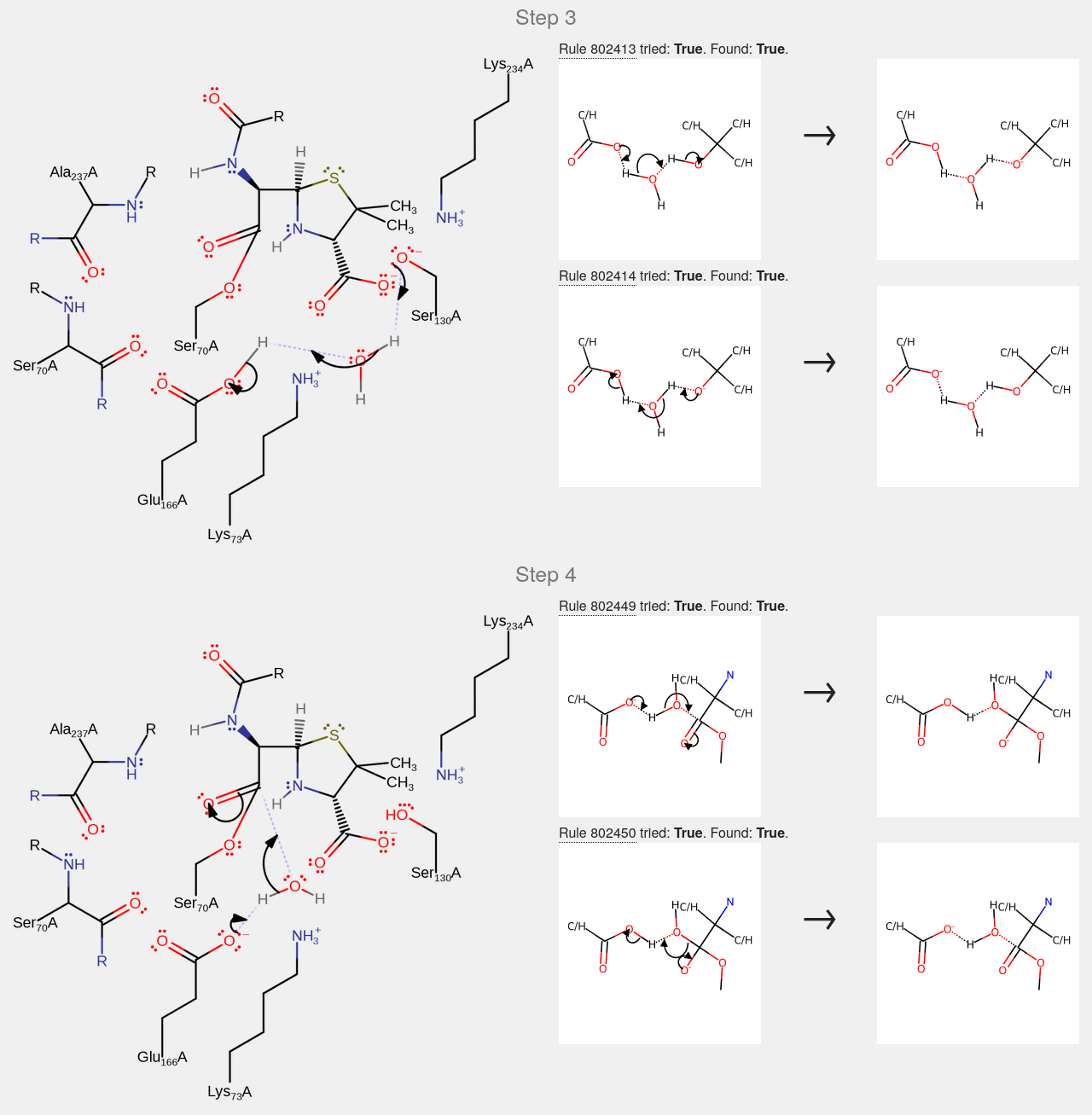

Figure SI 8 – Steps 3 and 4 of the mechanism of β-Lactamase A as annotated in the M-CSA. Rules extracted from each step are shown on the right as well as an indication of they were used in the prediction and if they were matched to any configuration.

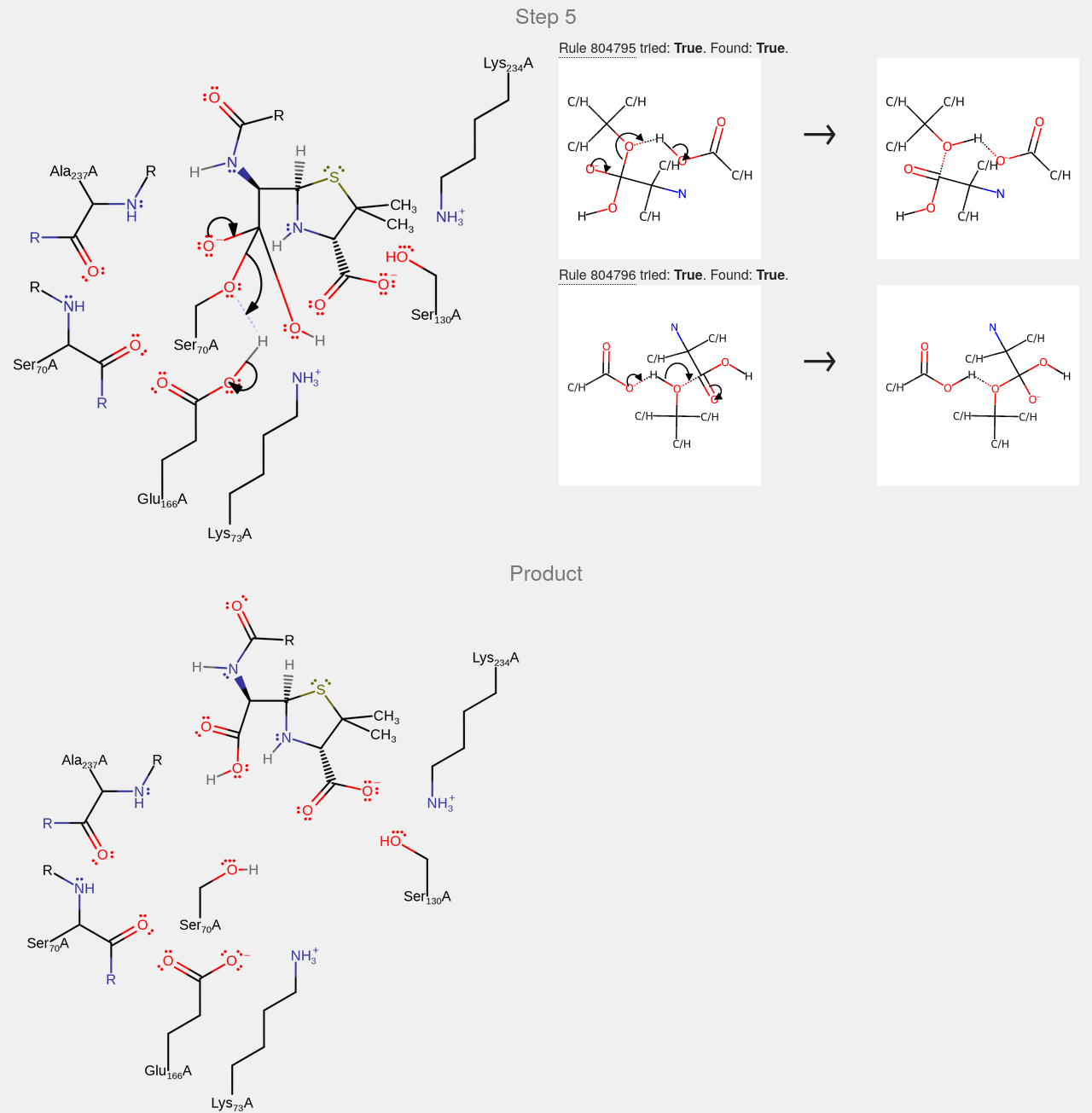

Figure SI 9 – Step 5 and products of the mechanism of β-Lactamase A as annotated in the M-CSA. Rules extracted from step 5 are shown on the right as well as an indication of they were used in the prediction and if they were matched to any configuration.

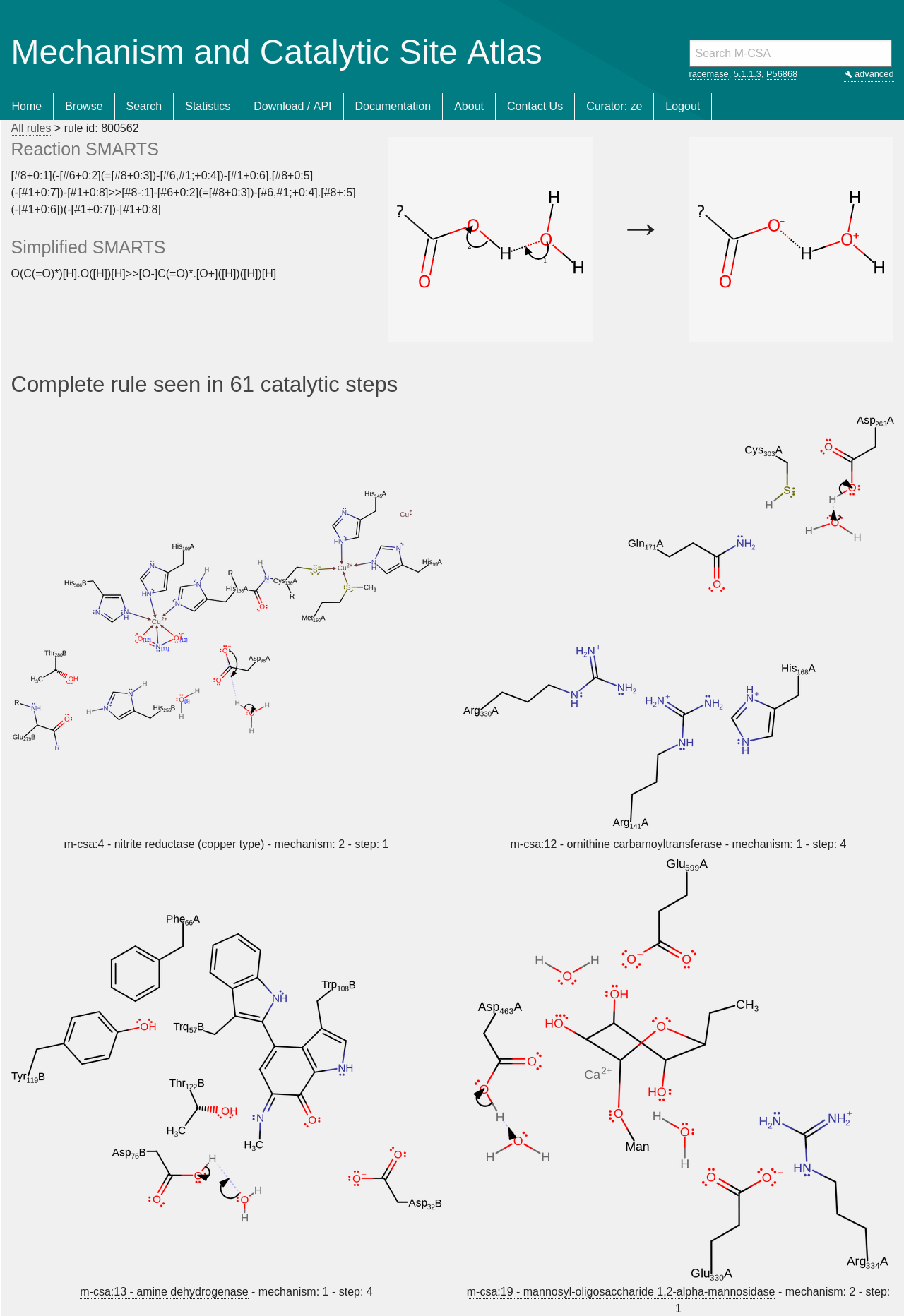

Figure SI 10 - Example of a rule page that shows the rule SMARTS pattern and a graphical representation of the rule. Below are linked all the catalytic steps where this rule is observed.
